# Evaluation of AlphaFold Antibody-Antigen Modeling with Implications for Improving Predictive Accuracy

**DOI:** 10.1101/2023.07.05.547832

**Authors:** Rui Yin, Brian G. Pierce

## Abstract

High resolution antibody-antigen structures provide critical insights into immune recognition and can inform therapeutic design. The challenges of experimental structural determination and the diversity of the immune repertoire underscore the necessity of accurate computational tools for modeling antibody-antigen complexes. Initial benchmarking showed that despite overall success in modeling protein-protein complexes, AlphaFold and AlphaFold-Multimer have limited success in modeling antibody-antigen interactions. In this study, we performed a thorough analysis of AlphaFold’s antibody-antigen modeling performance on 429 nonredundant antibody-antigen complex structures, identifying useful confidence metrics for predicting model quality, and features of complexes associated with improved modeling success. We show the importance of bound-like component modeling in complex assembly accuracy, and that the current version of AlphaFold improves near-native modeling success to over 30%, versus approximately 20% for a previous version. With this improved success, AlphaFold can generate accurate antibody-antigen models in many cases, while additional training may further improve its performance.

## Introduction

Antibodies are a key component of the immune system, defending the host from viruses and other pathogens through specific recognition of protein and non-protein antigens. Typically, antibodies engage their antigenic targets using the hypervariable complementarity determining region (CDR) loops within the variable domain^1^, which are stabilized by the β-sandwich structure of the framework region^2^. Despite sharing a conserved immunoglobulin structure, antibodies collectively exhibit a remarkable ability to recognize and bind to a wide array of antigens with high specificity. The highly specific and diverse nature of antibody-antigen interactions makes antibodies highly useful as therapeutics as well as a consideration in vaccine development efforts^3–6^.

High resolution structures of antibody-antigen complexes have refined our knowledge of immunity^7^, revealed molecular basis of antibody recognition of viral epitopes^8–10^, and guided the effective design of antibodies^11, 12^ and immunogens^13^. However, due to the challenges of experimental structure determination, resource and time constraints, as well as the highly diverse nature of the immune repertoire^14, 15^, experimental characterization of most antibody-antigen complex structures is impractical. Therefore, computational tools have been developed and applied to bridge this gap. General protein-protein docking methods have been applied to model antibody-antigen complex structures with limited success^16^, due in part to the need to account for the mobility of key CDR loops, as well as the size of certain antigens. To address this, algorithms have been developed specifically for antibody-antigen complex modeling^17–20^. However, accurate structural prediction of antibody-antigen complexes remains a challenge^16, 21^.

Recently, the scientific community saw a major breakthrough with AlphaFold (v.2.0), which uses an end-to-end deep neural network designed to predict protein structures from sequence^22^. AlphaFold iteratively infers and refines pairwise residue-residue evolutionary and geometric information from multiple sequence alignments (MSA) and has achieved unprecedented success in protein structure prediction^22, 23^. Its capabilities were expanded by the development of AlphaFold-Multimer^24^ (released in AlphaFold v.2.1), an updated implementation of AlphaFold that was designed to predict protein-protein complex structures. The overall architecture of AlphaFold-Multimer is similar to the previous version of AlphaFold, with changes including cross-chain MSA pairing, adjusted loss functions, and training on protein-protein interface residues. Previously our benchmarking revealed that, while generally successful in protein-protein complex structure prediction, AlphaFold was less successful in modeling antibody-antigen complexes, and adaptive immune recognition in general^25^. This lack of success in antibody-antigen structure prediction was also noted by the developers of AlphaFold-Multimer^24^. Yet, some high accuracy antibody-antigen complex models were generated by AlphaFold^25^, which shows potential for success of the “fold-and-dock” approach for antibody-antigen structure prediction. Given these initial limited reports, there is a need for extensive benchmarking and analysis of AlphaFold performance on these challenging and important targets.

Here we report a comprehensive benchmarking of AlphaFold performance for antibody-antigen complex structure modeling. With a dataset of over 400 high resolution and non-redundant antibody-antigen complexes, we investigated factors contributing to the successes and failures of the modeling process, including geometric and biochemical features of the complexes, and MSA depth. To distinguish accurate predictions from incorrect ones, we analyzed the use of AlphaFold-generated scores, as well as an interface confidence metric derived from AlphaFold residue accuracy (pLDDT) scores^25^. Furthermore, detailed analysis was conducted on the impact of recycling iterations, as well as the use of custom template inputs to evaluate the impact of accurate subunit modeling on antibody-antigen complex modeling accuracy. Finally, we benchmarked a recently released version of AlphaFold (v.2.3) and compared its performance to that of the previous version (v.2.2). Our study presents a thorough analysis of AlphaFold’s ability to predict antibody-antigen complexes, yielding valuable insights for interpreting model accuracy, identifying obstacles in the modeling process, and highlighting potential areas for improvement.

## Results

### AlphaFold-Multimer antibody-antigen complex modeling accuracy

To perform a comprehensive and detailed assessment of AlphaFold’s ability to model antibody-antigen complexes, we assembled a set of over 400 nonredundant antibody-antigen complexes released after April 30, 2018 (**Table S1**). The date cutoff was selected to avoid overlap with the training set of the tested version of AlphaFold (v2.2.0, hereafter denoted as v2.2 for brevity). Nonredundancy and additional test case selection criteria are described in the Methods section. For efficiency, we only utilized the variable domains of the antibody sequences for modeling. The accuracy of antibody-antigen complex predictions was evaluated using Critical Assessment of Predicted Interactions (CAPRI) criteria^26^, which classify predictions as Incorrect, Acceptable, Medium, or High based on a combination of interface root-mean-square distance (I-RMSD) and ligand root-mean-square distance (L-RMSD) from the experimentally determined complex structure, as well as the fraction of experimentally observed interface residue contacts in the model.

AlphaFold generated Acceptable or higher accuracy models as top-ranked predictions for 25% of the 429 test cases for which models were generated (**Fig. 1a**). Medium or higher accuracy models, which we refer to as near-native predictions, were generated as top-ranked predictions for 18% of the cases, and High accuracy models were generated for 5% of the test cases. Success rates increased when all 25 predictions per complex were taken into consideration, leading to 37% of the cases achieving Acceptable or higher accuracy predictions, 22% achieving Medium or higher accuracy predictions and 6% achieving High accuracy predictions.

**Figure 1.**
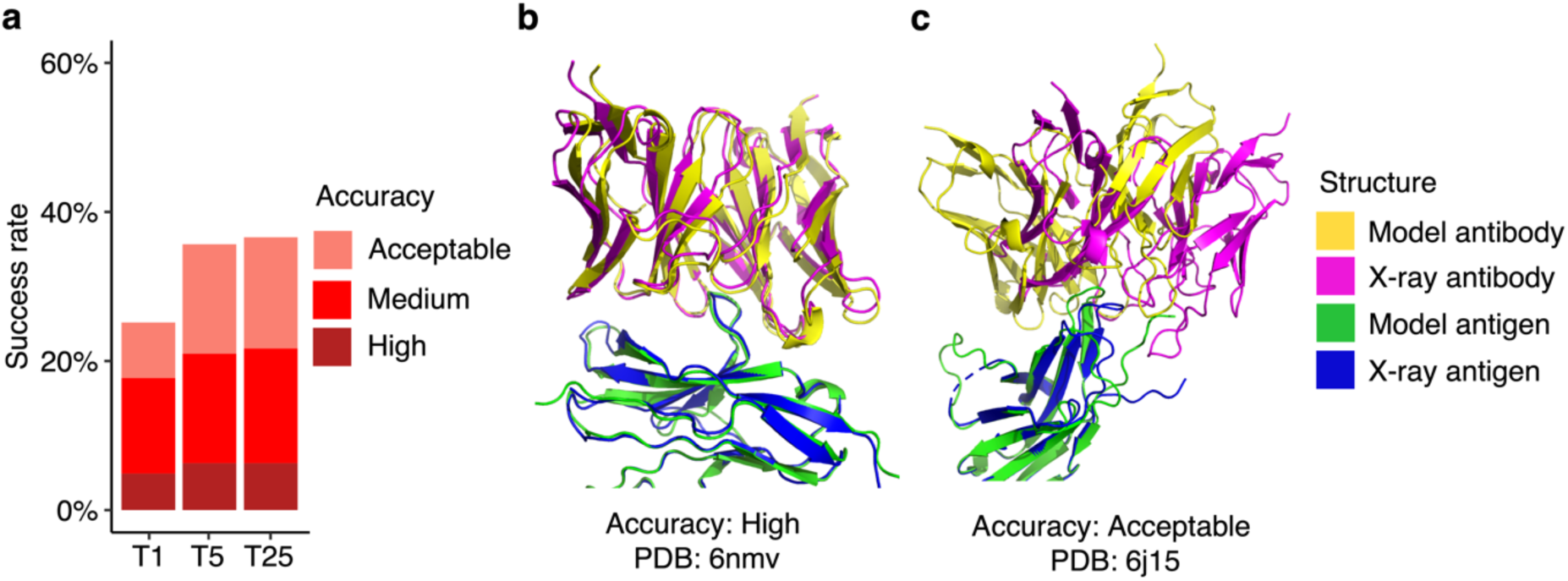
Antibody-antigen modeling success of AlphaFold. **a** Benchmarking of AlphaFold (v.2.2, AlphaFold-Multimer model) was performed on 429 antibody-antigen complexes. For each complex, 25 predictions were generated, and ranked by AlphaFold model confidence score. Antibody-antigen predictions were evaluated for complex modeling accuracy using CAPRI criteria for High, Medium, and Acceptable accuracy. The success rate was calculated based on the percentage of cases that had at least one model among their top N ranked predictions that met a specified level of CAPRI accuracy. Bars are colored by CAPRI accuracy level. **b** Example of a near-native prediction by AlphaFold, in comparison with the experimentally determined structure (PDB 6nmv; antibody/SIRP-alpha complex). This model has High CAPRI accuracy (I-RMSD=0.68 Å) and has the highest model confidence of all 25 predictions of this complex (model confidence=0.88). **c** An example of an Acceptable accuracy complex model from AlphaFold, in comparison with the experimentally determined structure (PDB: 6j15; antibody/PD-1 complex). This model has Acceptable CAPRI accuracy (I-RMSD=3.35 Å), and has the highest model confidence of all 25 predictions of this complex (model confidence=0.75). Complex structures in b and c are superposed by antigen with the model and the X-ray structure components colored separately as indicated on right.

Representative models generated by AlphaFold are shown in **Fig. 1b** (PDB code 6nmv; antibody/SIRP-alpha complex) and **Fig. 1c** (PDB code 6j15; antibody/PD-1 complex). Both models are top-ranked predictions for the respective complex. The model in **Fig. 1b** has High CAPRI accuracy, and an interface root-mean squared distance (I-RMSD) value of 0.68 Å, indicating a low level of structural deviation of this modeled antibody-antigen complex from the native complex. **Fig. 1c** shows an Acceptable CAPRI accuracy prediction with an I-RMSD of 3.55 Å. While the antibody engages the correct site of the antigen in this example, a deviation in positioning of the antibody on the antigen, with respect to the experimentally determined structure, is observed.

We additionally assessed antibody-antigen modeling accuracy of AlphaFold in ColabFold^27^. For fairness of the comparison with the full AlphaFold pipeline’s results, ColabFold was modified to generate 25 predictions per complex. ColabFold’s modeling success was similar to that of AlphaFold for 428 cases for which both algorithms were able to generate models, with slightly lower success observed for ColabFold (**Fig. S1**). The difference in success may be due to factors such as different MSAs or structural templates, as ColabFold and AlphaFold employ distinct approaches for building and pairing multiple sequence alignments, and utilize different sequence and template databases.

We observed higher success in modeling antibody-antigen complexes by AlphaFold here versus our previous benchmarking study, in which fewer than 10% of cases had top-ranked predictions with near-native accuracy^25^ (versus 18% here, as noted above). This difference is likely due to the newer version of AlphaFold used in this study (v2.2 vs. v2.1), which uses a retrained multimer model, as well as different sets of test cases, with the current study representing a substantial expansion over the cases used previously.

To compare antibody-antigen modeling performance with a previously developed docking approach, we utilized the global rigid-body docking algorithm, ZDOCK (version 3.0.2)^28^, with the IRAD (Integration of Residue- and Atom-based Potentials for Docking) reranking function which was developed to improve the ranking of near-native ZDOCK models^29^. Since many of the antibody-antigen complexes in our benchmark set do not have experimentally determined unbound antibody and/or antigen structures, AlphaFold was employed to generate unbound antibody and unbound antigen inputs for ZDOCK, using templates released on or before April 30, 2018. We selected the top-ranked prediction from AlphaFold as the input for ZDOCK, and only performed ZDOCK docking with unbound structures having a minimum average pLDDT score over 80, in order to exclude cases with likely low quality input modeled structures. In total, 390 complexes met the criteria for minimum average pLDDT score cutoff. Using ZDOCK with IRAD reranking, the majority of test cases did not yield highly accurate predictions (1% Medium or higher accuracy, **Fig. S2a**) as the top-ranked prediction. In contrast, AlphaFold-generated antibody-antigen complex models have a higher percentage of cases (19%) with Medium or higher accuracy top-ranked predictions within this set (**Fig. S2b**). The success rate increases when we consider top-25 predictions generated by ZDOCK (7% of complexes have Medium or higher accuracy predictions, **Fig. S2a**), yet the success is still lower than AlphaFold, which produced Medium or higher accuracy predictions for 23% of cases when all 25 predictions were considered. This ZDOCK success is similar to the unbound antibody-antigen docking success in Guest et al.^30^, although the difference in test cases, inputs (unbound versus modeled structures), and ZDOCK sampling and model scoring do not support a direct comparison of success rates. Challenges for ZDOCK likely responsible for the relatively low observed success rates include possible local inaccuracies in the unbound models, binding conformational changes (e.g. in CDR loops) even in the case of ideal unbound models, as well as modeled flexible protein terminal regions that are not resolved in some of the structures, which can lead to false positive binding sites for ZDOCK models.

### Antibody-antigen modeling accuracy determinants

To identify possible factors associated with modeling outcome, we analyzed properties of the native antibody-antigen complexes in relation to predictive modeling success. As glycans are not modeled by AlphaFold and antigen glycosylation can be an important component in antibody-antigen recognition (in some cases with glycans contacted directly by antibodies)^31^, the subset of complexes with antibody-antigen interface glycans in our set was identified to assess the impact of antigen glycosylation on modeling outcome. As some X-ray or cryo-EM structures used for analysis may lack resolved glycan atoms, or naturally occurring glycans can be removed enzymatically or via mutation to enable structural characterization, it is possible that some members of the non-interface glycan set (N=382) may have interface-proximal glycans in vivo. Our analysis showed that the presence of glycans at the native antibody-antigen interface is associated with lower modeling success (**Fig. 2a**). Among a total of 47 the cases with glycans at the interface, the top-ranked predictions of Medium accuracy were produced in only 9% of the cases, and no High accuracy top-ranked predictions were produced. In contrast, for cases not belonging to this category, 19% of them generated top-ranked predictions of Medium or higher accuracy. Thus, the lack of explicit consideration of glycans may reduce modeling accuracy for some antibody-antigen complexes.

**Figure 2.**
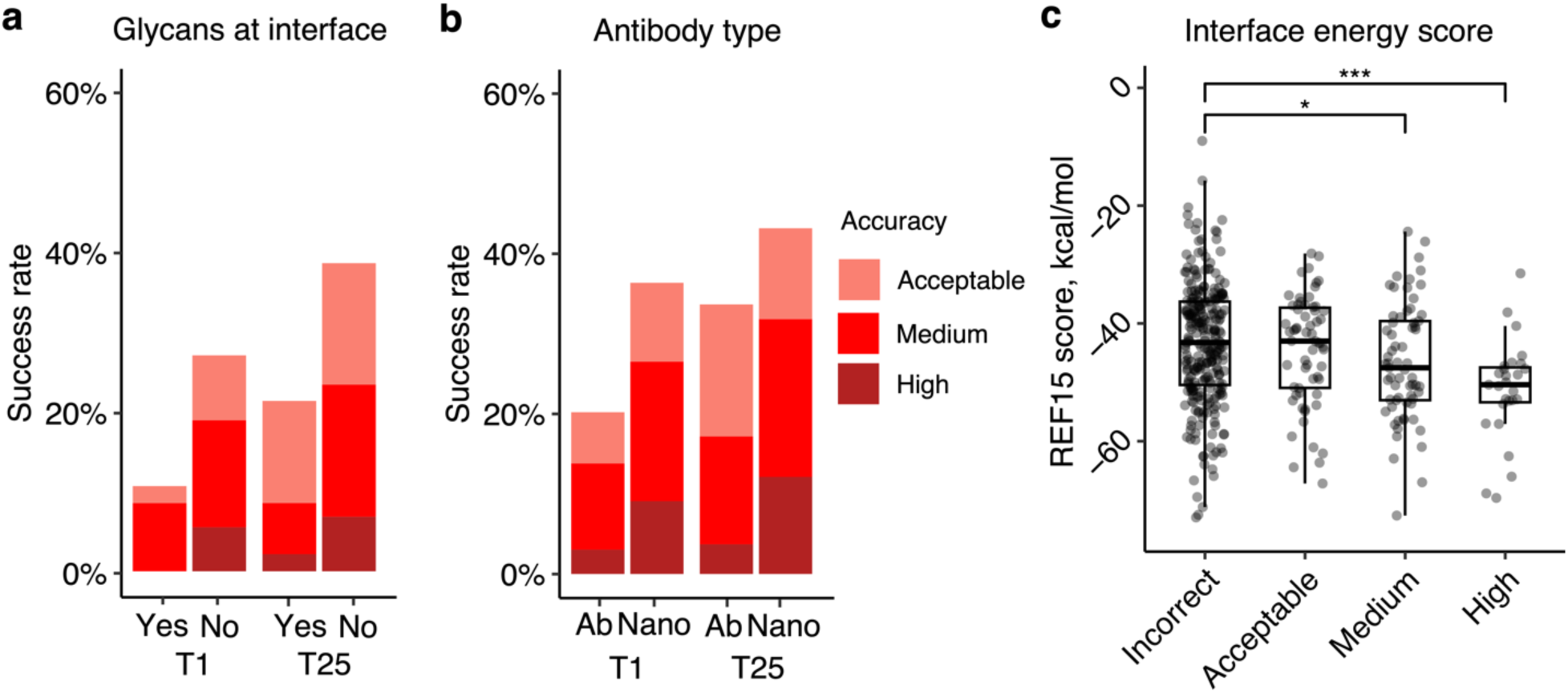
Properties associated with antibody-antigen modeling success. **a** Presence of glycans at the antibody-antigen interface. Glycans were identified through inspection of HETATM records residue classified as “saccharide” ligand, within 4.5 Å of from the antibody chain, in experimentally resolved structure coordinates. Complexes were classified as either “Yes” (N=47) or “No” (N=382) to indicate whether glycans were present or absent in the antibody-antigen interface. **b** Type of antibody in the complex. Complexes were classified as “Ab” (heavy-light antibody, N=297), or “Nano” (nanobody/VHH, N=132) based on antibody type. T1 and T25 denote AlphaFold modeling accuracy in top 1 (ranked by AlphaFold model confidence score) and in all 25 predictions of the complex. Bars were colored by CAPRI criteria. **c** Distribution of Interface energy score calculated by Rosetta InterfaceAnalyzer^62^ protocol (based on Rosetta REF15 energy function^60^) grouped by AlphaFold modeling accuracy. The modeling accuracy is defined as the highest CAPRI criteria prediction in the complex, considering all 25 predictions. Statistical significance values (Wilcoxon rank-sum test) were calculated between interface energy scores for sets of cases with Incorrect versus Medium and Incorrect versus High CAPRI accuracy predictions, as noted at top (* : p ≤ 0.05, *** : p ≤ 0.001).

We also investigated whether antibody-antigen complexes containing single-chain antibodies (or nanobodies) are more successfully modeled compared to the heavy-light chain only counterparts (**Fig. 2b**). For nanobody-antigen complexes (N=132), 27% of cases had Medium or higher accuracy top-ranked predictions, versus 14% of cases with Medium or higher accuracy top-ranked predictions for heavy-light chain antibody-antigen complexes (N=297). To understand the pronounced difference in modeling nanobody-antigen complexes versus antibody-antigen complexes, we investigated the difference in MSA depth of the two types of complexes. We hypothesized that the single-chain variable domains in nanobodies may simplify construction of cross-chain MSAs for nanobody-antigen complexes, as opposed to the more complex heavy-light chain antibodies. However, after analyzing the MSA depth, we found no statistically significant difference in the number of effective sequences (N_eff_, a measure of the effective sequence count in an MSA^22^) between the two types of complexes (**Fig. S3**). This suggests that other factors, such as fewer CDR loops and a smaller search space, may contribute to the observed difference in modeling success. Unlike heavy-light chain antibodies, which possess six CDR loops, the variable domain of nanobodies contains three loops only, thus it is possible that the lower complexity and size of the receptor component of the complex may play a role in the observed improved modeling performance for AlphaFold.

To investigate whether more energetically favorable antibody-antigen interfaces are more successfully predicted by AlphaFold, we compared antibody-antigen interface energy, computed from the bound complex structure using Rosetta^32^, with modeling success considering all 25 predictions of case (**Fig. 2c**). We found that more negative interface energies, indicative of more energetically favorable protein-protein interactions, are associated with higher AlphaFold modeling success. The difference in distribution of interface energy scores between complexes is statistically significant between Incorrect vs Medium accuracy prediction (p ≤ 0.05), and Incorrect vs High accuracy complexes (p ≤ 0.001), based on Wilcoxon rank-sum test.

While all complexes were modeled with antigen chains comprising the full epitope, a subset of cases did not include additional chains from the full antigen multimeric assembly (based on PDB “bioassembly”) in the modeling, due to computational limitations and for modeling efficiency. We examined whether these “partial antigen assembly” cases (N=51) were not as successfully modeled as the non-multimer or full assembly cases (N=378, **Fig. S4a**), due for instance to non-native surface regions that are normally buried in the full antigen assembly being engaged by antibodies in the models. Indeed, the partial antigen assembly cases exhibited lower modeling success versus the remainder of the cases. Additionally, consistent with our previous benchmarking study^25^, success analysis on the current set of antibody-antigen complexes shows that larger complexes are generally more difficult to predict (**Fig. S4b**).

We also examined the accuracy of six complementarity determining region (CDR) loops in the modeled complexes, computing CDR loop RMSDs in the models with respect to the corresponding CDR loop from the experimentally determined complex structure (**Fig. S5**).

Interestingly, the CDRH3 loop exhibited differences in accuracy across sets of models with different CAPRI accuracy levels, with median CDRH3 RMSD of 0.6 Å for High accuracy models, 1.2 Å for Medium accuracy models, 2.2 Å for Acceptable accuracy models, and 2.4 Å for Incorrect models, indicating that accurate prediction of CDRH3 loops is associated with near-native modeling accuracy in antibody-antigen complex prediction in AlphaFold. However, it should be noted that CDR loop accuracy (among other features of the modeled interfaces) can play a role in the CAPRI antibody-antigen complex accuracy assessments themselves, with High CAPRI accuracy models likely requiring relatively low CDRH3 RMSDs to closely reflect the native interface.

We also examined possible failures of antigen structure modeling as factors in antibody-antigen modeling success (**Fig. S6**). The analysis of the antigen accuracy in top-ranked complex predictions revealed that while only 12 complexes out of 429 complexes have antigen predictions with TM-score^33^ values below 0.7 with respect to the experimentally determined antigen, indicating a relatively lower level of structural similarity^34^, the majority of complex predictions included relatively accurate modeling of the antigen subunit. This suggests that most of the failed predictions can be primarily attributed to incorrect docking poses or local structural perturbations rather than inaccurate antigen subunit predictions.

### Model confidence score comparison

The reported success of model accuracy scores produced by AlphaFold^24, 25^ led us to evaluate the ability of those scores, or adaptations thereof, to discriminate between accurate vs. incorrect antibody-antigen predictions. We assessed AlphaFold’s model confidence score, which is a linear combination of pTM and ipTM^24^ scores, as well as interface pLDDT (I-pLDDT), which is based on residue-level confidence scores for antibody-antigen interface residues (4 Å distance cutoff), as used in previous studies^25, 35^, for discrimination of correct antibody-antigen models (**Fig. 3**). While both exhibited significant correlations with DockQ score^36^, which is a continuous measure of complex model accuracy, I-pLDDT was marginally superior (**Fig 3a-b**); this was also evident for comparison of the scores with CAPRI accuracy levels (**Fig. 3c-d**).

**Figure 3.**
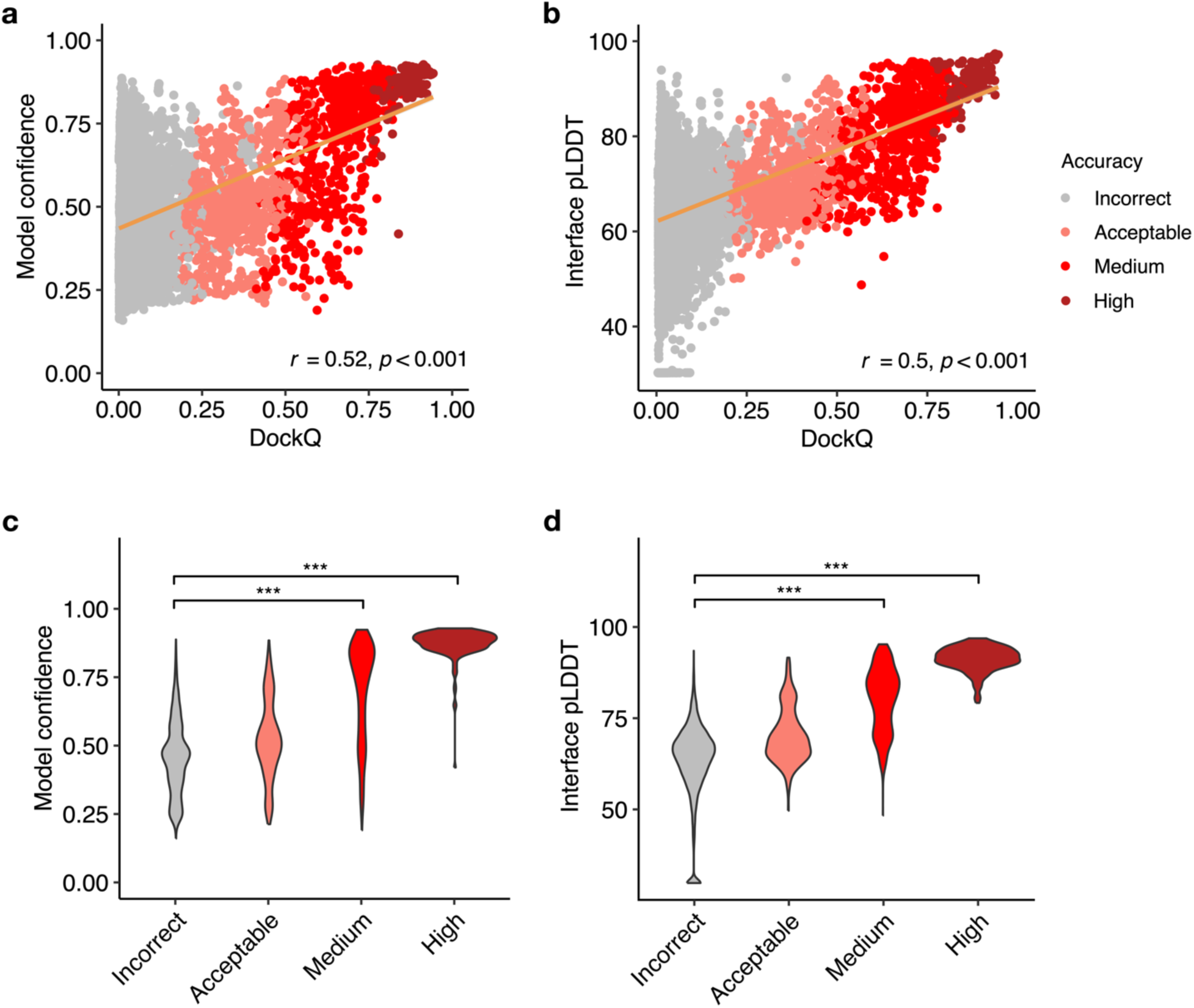
AlphaFold model confidence scores and model accuracy. Scatter plots compare **a** model confidence and **b** interface pLDDT score with model accuracy, with accuracy assessed by DockQ score. In the scatter plots, all 25 models representing 429 complexes are depicted as data points, with their colors indicating the model quality according to CAPRI criteria. The orange line represents the linear regression, and the lower right corner of the scatter plots displays the Pearson’s correlation coefficients and correlation p-values. Distribution of **c** model confidence and **d** interface pLDDT score, grouped by the CAPRI criteria of AlphaFold predictions. Interface pLDDT score is defined as the mean of pLDDT scores of residues within 4 Å of the antibody-antigen interface. Complexes without contacts within 4 Å of antibody-antigen interface is assigned a pLDDT score of 30. Statistical significance values (Wilcoxon rank-sum test) were calculated between model scores for sets of predictions with Incorrect versus Medium and Incorrect versus High CAPRI accuracy, as noted at top (*** : p ≤ 0.001).

I-pLDDT also provided outstanding discrimination between Incorrect vs. Medium or higher accuracy models based on receiver operating characteristic area under the curve (AUC) metrics (AUC=0.93), which is higher than that of the model confidence (AUC=0.88; **Table 1**). We also tested the individual components of the model confidence scores (pTM and ipTM) (**Fig. S7**), which did not yield improved correlations with DockQ scores versus model confidence. When excluding data points without side-chain contacts within 4 Å across the antibody-antigen interface (for which I-pLDDT was set to an arbitrary minimum value in **Fig. 3b** and in the corresponding correlation calculation), the correlation between the interface pLDDT and DockQ increased to *r*=0.57 (**Fig. S8a**), which demonstrates a more significant difference compared to the correlation between the model confidence and DockQ (*r*=0.53, **Fig S8b**).

**Table 1.** Area under the ROC curve (AUC) value for protein model quality classes as a function of different scoring metrics.

| Score<sup>a</sup> | Binary classification<sup>b</sup> |  | Multi-class classification<sup>b</sup> |
| --- | --- | --- | --- |
|  | Incorrect vs. High | Incorrect vs. Medium and High |  |
| interface pLDDT | 1.00 | 0.93 | 0.88 |
| model confidence | 0.99 | 0.88 | 0.85 |
| ipTM | 0.99 | 0.87 | 0.85 |
| pTM | 0.99 | 0.89 | 0.85 |

<sup>a</sup> Scoring methods. “interface pLDDT”: average pLDDT of antibody-antigen interface residues within 4 Å. Models without antibody-antigen interface contacts were assigned an interface pLDDT value of 30.
<sup>b</sup> The ROC values of binary classification and multi-class classification were calculated using the R pROC<sup>58</sup> and multiROC<sup>59</sup> packages. Protein model quality class is assigned as per CAPRI criteria, into Incorrect (n=9,112), Acceptable (n=773), Medium (n=684) and High (n=156) classes.

One advantage of I-pLDDT over ipTM and model confidence (which primarily consists of ipTM) is that it is specifically focused on the antibody-antigen interface, whereas ipTM is calculated across all inter-chain interfaces of complex models, including heavy-light and multiple antigen chains, thus the latter scores may be influenced by less relevant elements of the complex. Overall, these results support the use of I-pLDDT as a primary metric in assessing the quality of AlphaFold antibody-antigen models.

### Progressive improvements over recycling iterations

Recycling is a critical component of the AlphaFold algorithm^22, 24^, wherein each model is input back to the system for further optimization. To improve our understanding of the impact of recycling iterations on AlphaFold modeling of antibody-antigen complexes, we modified the AlphaFold pipeline in ColabFold. ColabFold was preferable to utilize in this context versus the default AlphaFold pipeline due to its speed, in order to enable output and analysis of the antibody-antigen complex predictions at each recycling iteration. Our analysis demonstrates an increase in model accuracy as recycling iterations progress (**Fig. 4a**). In fact, approximately 50% of predictions of Medium or higher accuracy after the third recycle iteration were Incorrect models before recycling iterations (**Fig. S9**).

**Figure 4.**
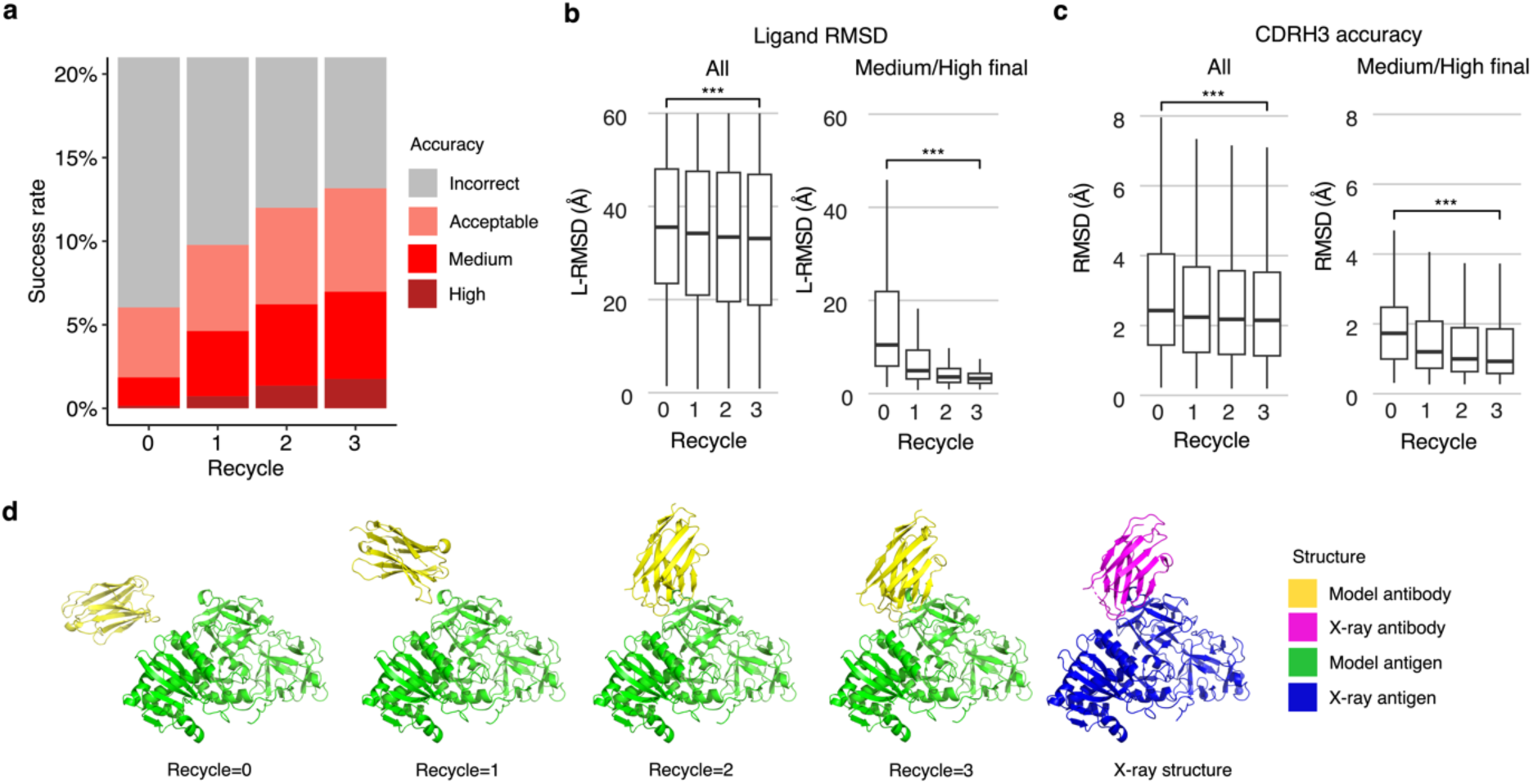
Analysis of antibody-antigen predictions across recycling iterations. **a** The accuracy of antibody-antigen complex predictions across up to three recycling iterations. Complex prediction accuracy across recycling iterations (up to three recycles, denoted by the x-axis). Success rate is defined as the proportion of predictions of specific level of CAPRI criteria in a total of 25 prediction per complex, 428 complexes total, at the given recycle. Recycle=0 denotes the state of the prediction before recycling iterations begin. **b** Distribution of the ligand RMSD (L-RMSD, Å) of antibody-antigen prediction at each recycling iteration (denoted by the x-axis), of all predictions (25 predictions * 428 complexes, left panel) or a subset of predictions of Medium or High CAPRI accuracy at recycle=3 (109 predictions, right panel). **c** Distribution of the CDRH3 accuracy of antibody-antigen prediction at each recycling iteration (denoted by the x-axis), of all or a subset of predictions of Medium or High CAPRI accuracy at recycle=3. CDRH3 accuracy is defined as the change in RMSD of the CDRH3 region, when superposing the predicted antibody (in the antibody-antigen complex prediction) onto the experimentally resolved antibody (in the antibody-antigen complex) using the antibody framework region. Statistical significance values (Wilcoxon rank-sum test) were calculated between RMSD values for sets of predictions at the outset of recycling iterations (recycle=0) vs at recycle=3, as noted at top (***p ≤ 0.001). **d** Example of a prediction across recycling iterations (PDB 7kd2; nanobody/Ricin complex). This prediction’s CAPRI accuracy level across recycles was Incorrect at recycle=0 (I-RMSD=17.98 Å), Incorrect at recycle=1 (I-RMSD=10.90 Å), Acceptable at recycle=2 (I-RMSD=2.52 Å), and Medium at recycle=3 (I-RMSD=1.45 Å). The CDRH3 RMSD of the prediction across recycling iteration 0, 1, 2, and 3 was 1.39 Å, 1.19 Å, 1.27 Å, and 1.17 Å, respectively. The L-RMSD of the prediction across recycling iteration 0, 1, 2, 3 was 49.95 Å, 24.68 Å, 5.42 Å, 4.25 Å, respectively. Antibody and antigen chains of the predictions and x-ray structure were colored as indicated. Predictions were generated with ColabFold due to its faster model generation speed, compared to AlphaFold.

Next, we analyzed specific changes in antibody-antigen model across recycling iterations, identifying notably enhanced features and those that are unchanged. Features that were improved highlights the strength of AlphaFold, whereas the lack of improvement may highlight areas of difficulty or suggest that these features were already optimal at the start and did not require further refinement. We analyzed both the accuracy of antibody positioning on the antigen and the quality of the highly variable CDR loop of the antibody. Given the high variability in CDRH3 RMSD (**Fig. S4**), compared to the RMSD of other CDR loops, we focused our analysis of CDR loops on the CDRH3. Considering all predictions, we observed a marginal yet significant improvement in both the antibody-antigen binding conformation as measured by ligand RMSD (L-RMSD) (**Fig. 4b**, left panel) and the CDRH3 loop accuracy (**Fig. 4c**, left panel). Upon examining the subset of cases with Medium or higher accuracy at recycle 3, we observed that the antibody-antigen binding conformation score L-RMSD exhibited a pronounced and significant improvement (**Fig. 4b**, right panel), while the improvement in CDRH3 loop RMSD was significant but not as pronounced (**Fig. 4c**, right panel), indicating that for models to attain high accuracy at the end of the recycling iteration, it is helpful for AlphaFold to accurately predict the CDRH3 loop relatively accurately before recycling iterations begin.

The capability of AlphaFold to perform rigid-body protein movements over recycling iterations, is shown in **Fig. 4d** (nanobody/Ricin complex). This prediction was of Incorrect accuracy before recycling iterations and was improved to a Medium accuracy prediction at recycle 3. Over recycling iterations, the L-RMSD of this prediction exhibited a substantial degree of improvement, from 49.95 Å before recycling, to 4.25 Å at recycle 3. Unlike L-RMSD, the CDRH3 loop of this prediction was accurately predicted (CDRH3 RMSD=1.39 Å) before the recycling iterations.

The importance of CDRH3 loop accuracy for complex modeling success was further explored by the analysis of CDRH3 loop conformations of modeled unbound structures. Unbound antibody structures were generated with AlphaFold with a template date cutoff of April 30, 2018, and the CDR loops of the unbound antibody models were compared to those of the antibodies in the antibody-antigen complexes. The RMSD between CDR loops of the unbound models and the antibody in the bound is compared against the complex modeling success of top-ranked antibody-antigen models generated by AlphaFold in **Fig. S10**. Although the relatively small numbers of High accuracy cases limits this comparison, the accuracy of the CDRH3 modeling in unbound antibody structures for High antibody-antigen models was found to be significantly higher than that of the Incorrect accuracy models (p ≤ 0.05), suggesting that antibodies with unbound models that more closely resemble the bound loop conformation are likely to be more accurately modeled in the form of antibody-antigen complexes.

### Accurate modeling of subunits in bound-like form enables higher success

To better understand the factors that can enhance the success rate of the AlphaFold antibody-antigen modeling, we utilized native antibody-antigen chains as templates within the AlphaFold modeling pipeline, to gauge whether AlphaFold can better assemble the complex structures given the bound subunit chains. Modifications were made to the AlphaFold pipeline to optionally input specific selected PDB templates for each chain. To test performance, we randomly selected 100 cases from the full antibody-antigen benchmark that do not have observed glycans at the antibody-antigen interface and do not belong to the partial antigen assembly category, due to observed change in performance for those sets of cases (**Fig. 2**, **Fig. S4**). On this subset of 100 cases, the use of default templates identified from the AlphaFold pipeline resulted in 18% success in generating near-native (Medium or High accuracy) top-ranked predictions (**Fig. 5a**), which is similar to the performance on the full benchmark (**Fig. 1a**). A substantial improvement in accuracy was observed when experimentally determined antibody-antigen chains were used as individual chain templates, in which case the success in generating near-native top-ranked predictions was 52% (**Fig. 5b**). Analysis of the top-ranked prediction success determinants shows that distribution of interface energy score (**Fig. S11a**) and change in solvent-accessible surface area (ΔSASA) for hydrophobic part of the antibody-antigen interface (**Fig. S11b**) are significantly different (p ≤ 0.01) between complexes that have Incorrect versus High accuracy top-ranked predictions, indicating that despite using bound template structures, AlphaFold has difficulty predicting the complex structure for antibody-antigen interactions with less favorable computed interface energies and with smaller hydrophobic interface area.

**Figure 5.**
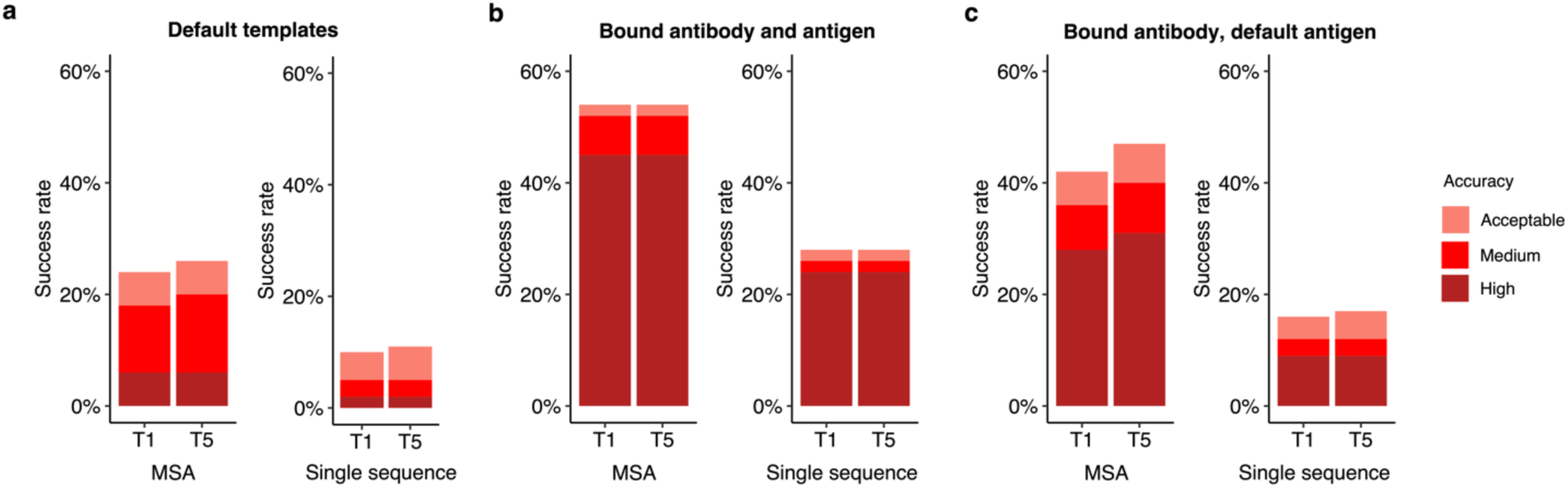
Improved subunit modeling enhances antibody-antigen complex modeling success. Antibody-antigen modeling success of AlphaFold by utilizing **a** templates identified through the default template search protocol, **b** bound antibody and antigen chains as templates, **c** bound antibody and default antigen chains (identified by the default search protocol) as templates. Benchmarking was performed on a total of 100 antibody-antigen complexes. The success rate was calculated based on the percentage of cases that had at least one model among their top N predictions that met a specified level of CAPRI accuracy. Bars are colored by CAPRI accuracy criteria.

For comparison of predictive success with bound component inputs, we employed ZDOCK^28^ with IRAD^29^ scoring to perform global rigid-body docking and ranking with the bound antibody and antigen structures (with randomized initial orientations). This approach led to a relatively high proportion (62%) of cases having top-ranked predictions with Medium or higher accuracy (**Fig. S12a**), indicating that both traditional docking as well as AlphaFold can successfully assemble over half, antibody-antigen complex structures with bound components, while still unable to assemble a sizable fraction of the them.

Compared to using all experimentally determined chains as templates, utilizing only certain bound chains as templates resulted in decreased model accuracy. Specifically, with default antigen templates, and experimentally determined antibody heavy and light chains provided as templates, 36% of top-ranked predictions were of Medium or higher accuracy (**Fig. 5c**). A similar near-native success rate (35%) was observed when using only bound antibody heavy chains as templates (**Fig. S12b**), while light chain and antigen bound chain templates led to 28% of cases with near-native top-ranked predictions (**Fig. S12c**). Overall, the resulting success rates of these bound chain template scenarios are higher than those obtained using the default templates, in which case 18% of the complexes have Medium or higher accuracy top-ranked predictions. While not reflective of actual predictive modeling scenarios due to the use of bound structure information, these results indicate the potential impact of accurate and bound-like subunit modeling, as well as its theoretical maximal effect, for AlphaFold antibody-antigen modeling success.

Inspired by the findings, we tested using modeled subunit structures as templates in antibody-antigen modeling, as we hypothesized that if these models are more accurate than the default pipeline’s selected template structures, they could potentially improve AlphaFold’s performance over the default pipeline and templates. However, this yielded lower success than observed when using default templates (12%, versus 18% for near-native modeling success) (**Fig. S12d**), showing that inaccuracies or deviations from the bound components for the unbound models led to far different behavior than when using actual bound components.

### MSA provides important information for accurate prediction of complexes

We also evaluated the performance of AlphaFold without MSAs, to test the impact on complex assembly when subunit structures are known (thus MSA would not in principle be needed for subunit structure modeling), given the likely lack of direct co-evolutionary information present in antibody-antigen MSAs. The removal of MSAs was implemented through modifications to the AlphaFold pipeline, as noted in the Methods. Our results indicated a notable decrease in accuracy when MSA was disabled, as compared to the with-MSA counterparts (**Fig. 5** and **Fig. S12b**,**c**). This prompted us to investigate the possible association between the depth of MSA and the modeling outcome by AlphaFold.

We investigated the impact of MSA depth on modeling success by the full AlphaFold protocol, grouping the complexes by prediction accuracy and comparing distributions of MSA depth (N_eff_) (**Fig. 6**). The distribution of N_eff_ was found to be statistically significant between Incorrect and Medium accuracy classes (p ≤ 0.01), and between Incorrect and High accuracy classes (p ≤ 0.01). We also compared the docking model quality (DockQ score) for all cases when binned by MSA depth levels (**Fig. S13**). A slight trend was observed indicating that a greater MSA depth is associated with higher DockQ scores (higher model accuracies), suggesting that compared to a shallow MSA, predictions with a deeper MSA are more likely to be of higher accuracy. Thus, it is possible that increasing MSA depth, particularly for antibody-antigen complexes with very shallow MSAs, could lead to some improvement in overall modeling performance.

**Figure 6.**
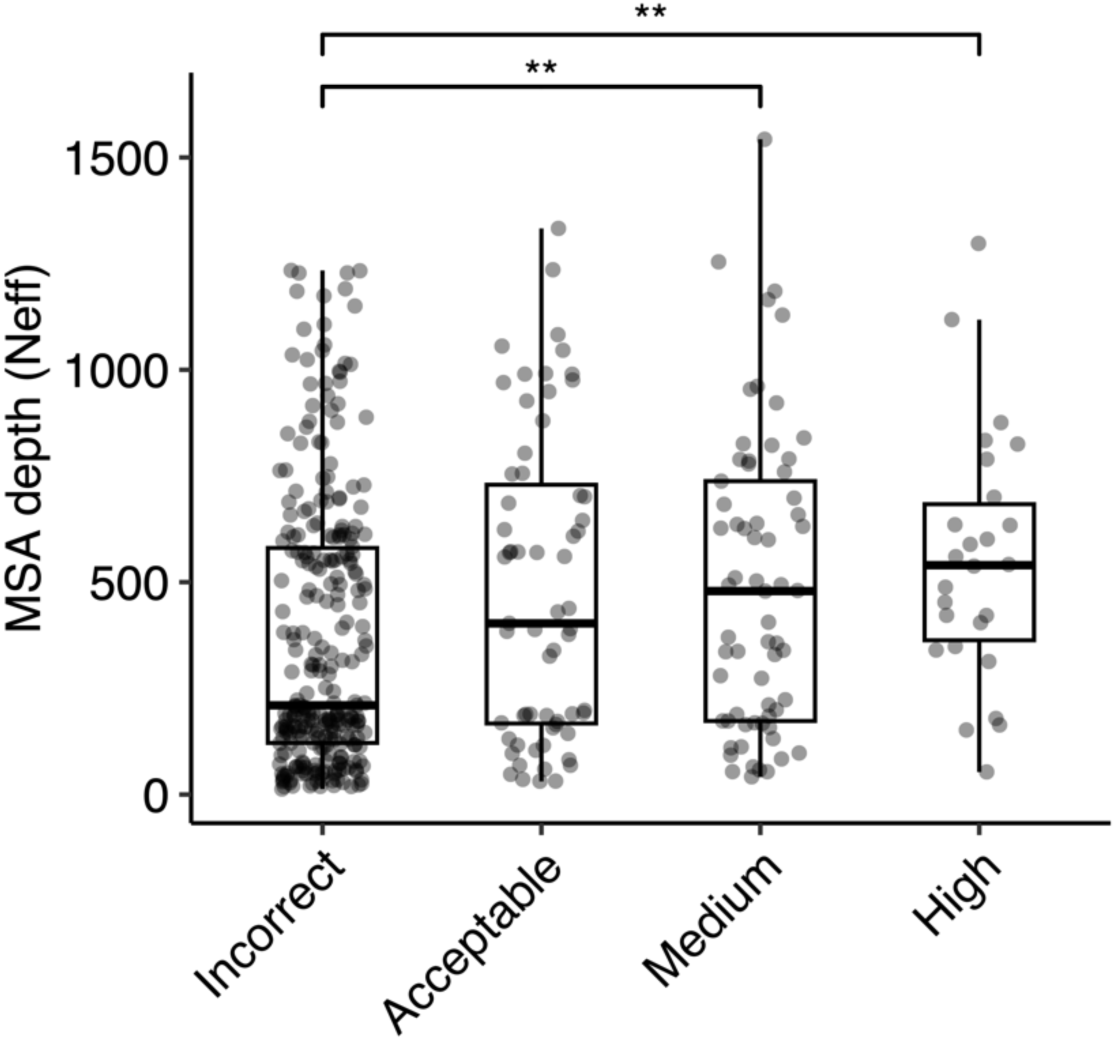
Comparison of MSA depth and modeling success. The distribution of MSA depth (number of effective sequences, N_eff_), calculated using CD-Hit^56^ with an identity cutoff of 80%, is shown for antibody-antigen complexes grouped by AlphaFold modeling accuracy. The modeling accuracy is defined as the highest CAPRI criteria prediction in the complex, considering all 25 predictions. Numbers of data points in Incorrect, Acceptable, Medium and High categories are 274, 63, 65 and 26. Statistical significance values (Wilcoxon rank-sum test) were calculated between interface energy scores for sets of cases with Incorrect versus Medium and Incorrect versus High CAPRI accuracy predictions, as noted at top (**p ≤ 0.01).

### Modeling accuracy of AlphaFold v.2.3.0

Recently, an updated version of AlphaFold (v.2.3.0, hereafter denoted as v.2.3) was released, with modifications to the pipeline and deep learning model^37^. Compared with the previous version, this version was trained on PDB structures released until September 30, 2021, resulting in a 30% increase in training data. This version also increased the maximum number of recycles, from 3 recycles in v.2.2 to 20 recycles in v.2.3, with early stopping, and utilized larger interface regions (crops) and more chains during training. To benchmark its performance, we assembled a test set of 41 nonredundant antibody-antigen complexes released after the September 30, 2021 training date (**Table S1**). On a total of 39 cases for which models were successfully produced by both versions of AlphaFold, v.2.3 generated Medium or higher accuracy models as top-ranked predictions for 36% of the test cases, notably higher than the 23% generated by v.2.2 (**Fig. 7**), with no significant difference in antibody CDR loop accuracy.

**Figure 7.**
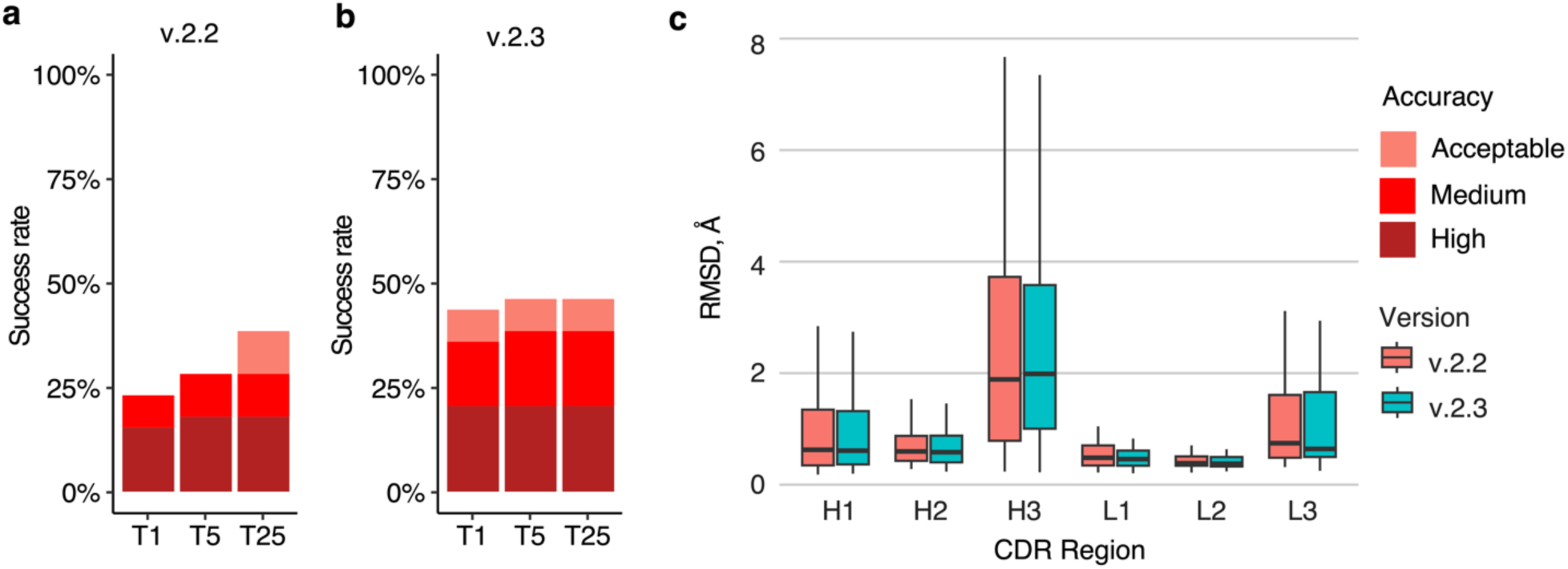
Antibody-antigen modeling success by AlphaFold v.2.3. Modeling success of **a** AlphaFold v.2.2 and **b** AlphaFold v.2.3 on 39 antibody-antigen complexes. The success rate was calculated based on the percentage of cases that had at least one model among their top N predictions that met a specified level of CAPRI accuracy. Bars were colored as per CAPRI accuracy criteria. **c.** Distribution of the CDR loop prediction accuracy of AlphaFold v.2.2 (denoted by “salmon” color) vs v.2.3 (denoted by “cyan” color). CDR loop accuracy is defined as the change in RMSD of the CDR regions, when superposing the predicted antibody (in the antibody-antigen complex prediction) onto the experimentally resolved antibody (in the antibody-antigen complex) using the antibody framework region.

Compared to v.2.2, v.2.3 predictions were produced with a higher number of recycles, with 95% of v.2.3 predictions generated using more than three recycles. To further investigate this observation, we reduced the maximum number of recycles in v.2.3 from 20 to 5 (**Fig. S14a**) or 3 (**Fig. S14b**). Compared to the default setting with a maximum number of recycles of 20, the antibody-antigen complex prediction success remained identical for generating near-native top-ranked predictions, regardless of the recycle limit. These findings suggest that the observed difference in the number of recycles between v.2.3 and v.2.2 is not the main factor contributing to the increased success, and that the updated and expanded training of the deep learning model training may be responsible.

## Discussion

Using a set of over 400 nonredundant antibody-antigen complexes, we benchmarked and evaluated AlphaFold’s ability to model antibody-antigen complexes. On this set, we observed a limited yet higher success in the prediction of antibody-antigen structures by AlphaFold, compared to our previous benchmarking that used an older AlphaFold version accessed via ColabFold, and was based on a limited set of 100 antibody-antigen cases^38^. Analyses of factors that could influence the prediction outcome showed that AlphaFold was less able to accurately predict antibody-antigen structures with glycans at the antibody-antigen interface, which highlights AlphaFold’s limitation in handling complexes with post-translational modifications. We also found that AlphaFold is more successful at modeling nanobody-antigen complexes and has difficulty predicting the structure of larger antibody-antigen complexes. An analysis of prediction accuracy at each recycling iteration, as well as the bound antibody-antigen template tests shows the importance of accurate subunit modeling for success in predicting the antibody-antigen complex. Relatedly, the ability to accurately predict CDRH3 loops is important for overall docking success.

Our benchmarking also shows that the latest version of AlphaFold (v.2.3) exhibits improved success in predicting antibody-antigen structures versus the previous AlphaFold version (v.2.2), likely due at least in part to the model training on an updated and expanded set of complex structures from the PDB^39^. It is possible that success can be improved further through additional optimization or other adaptations of the AlphaFold framework or model. Recent work has shown that higher structural diversity in AlphaFold predictions can be achieved by via enabling dropout^27, 36, 40^. When this technique is coupled with enhanced sampling, the quality of the generated predictions can be improved, as recently described^40^, while additional strategies that have been explored for protein conformational sampling in AlphaFold, as noted in a recent review^41^, could likewise be tested for antibody-antigen complexes. The observed strong association between antibody-antigen model accuracies and multiple confidence scores indicates that such increased sampling may be a worthwhile approach. Another potential avenue for elevating the accuracy of AlphaFold predictions is demonstrated by the recent development of fully trainable AlphaFold implementations^42–44^, which enable researchers to adapt and refine the model to specific datasets or domains of interest, opening up new possibilities for customization and optimization of the AlphaFold network.

Despite the lack of explicit coevolutionary signal, our data show that the inclusion of diverse sequence information in MSAs is helpful for maintaining AlphaFold’s modeling success of antibody-antigen complexes. As such, curation or optimization of MSAs could be another avenue for improving the accuracy of AlphaFold predictions. Previous work showed that AlphaFold prediction of protein-protein complexes can be augmented with improved MSA cross-chain pairing^35^, while others have developed alternative MSA methods such as DeepMSA2^45^, which was part of a successful pipeline in a recent CASP/CAPRI complex structure prediction round^46^. Recent work leveraging protein language models shows promise in constructing diversified and informative MSAs for enhancing accuracy in AlphaFold protein complex prediction^47^, while it may be possible to replace or augment the MSA in AlphaFold with language model representations, possibly building on recent language models developed for antibodies such as AntiBERTy^48^.

Our results also demonstrate that accurate subunit prediction is associated with higher antibody-antigen complex prediction success. Recent work has shown improved accuracy in antibody prediction, particularly in the context of CDR loops, leveraging elements of AlphaFold architecture, especially the structure module, with modifications^49, 50^. Incorporating such advances into the prediction pipeline may enable the prediction of more accurate antibody-antigen complexes.

While it is possible or even likely that antibody-antigen modeling success may ultimately be improved in AlphaFold or related deep learning frameworks, the current success of AlphaFold and version 2.3 in particular, in conjunction with the observed confidence scoring accuracy, indicates that AlphaFold may potentially be of practical use to researchers in modeling this important and challenging class of complexes, and can complement or assist experimental structural determination methods.

## Methods

### Antibody-antigen benchmark assembly

We assembled two nonredundant sets of high resolution structures to benchmark AlphaFold, following the general protocol that we described previously^25^. To obtain an initial list of antibody-antigen complexes from the PDB, we downloaded the full SAbDab antibody structure dataset in January 2022. The antibody-antigen complex dataset for AlphaFold v2.2 benchmarking was assembled using the following criteria: 1) structure resolution <= 3.0 Å, 2) protein antigen in the structure (based on SAbDab annotation), and 3) nonredundant with antibody-antigen complexes with structural resolution <= 9.0 Å released before April 30, 2018 (AlphaFold v2.2 training sample cutoff date) based on sequence criteria. Sequence criteria for nonredundancy are: 1) heavy chain variable domain sequence ID < 90% and full variable domain sequence ID < 90%, or 2) no match between antigen chain sequences (no hit detected using BLAST^51^ with default parameters). Pairwise sequence alignments were performed using the “blastp” executable in the BLAST suite^51^. Structural nonredundancy criteria were then applied to the set. We removed antibody-antigen structures with < 5 Å heavy chain Cα atom RMSD, after superposition of antigens using the FAST structure alignment program^52^, and > 70 % identity between heavy chain variable domain, light chain variable domain, or concatenated CDR loop sequences. To avoid modeling antigen chains with large regions that are not resolved in the experimentally determined structures, we additionally removed structures with PDB “seqres” file sequence annotation and resolved region sequence length difference > 70%, or sequence length difference > 35% and resolved antigen length > 500 aa. We also removed non-canonical antibody-antigen complex cases (e.g. with antibody-tetramerization, dimeric sdAb, or constant domain binding), and we removed cases with incomplete antigen chain annotations by SAbDab, identified through manual inspection of the PDB bioassembly structure.

To benchmark AlphaFold v.2.3, we identified a subset of 41 antibody-antigen complexes within the v.2.2 benchmarking set. These antibody-antigen complexes were released after September 30, 2021, and are not redundant with structures released before that date based on the sequence criteria detailed above. AlphaFold v.2.2 and v.2.3 generated models for 39 out of 41 of those complexes without runtime errors.

The AlphaFold v2.2 and v2.3 benchmarking cases are shown in **Table S1**.

### AlphaFold antibody-antigen modeling

Sequences input to AlphaFold were obtained from the PDB “seqres” file. Antibody sequences were processed by ANARCI to remove non-variable domain sequence regions. We downloaded and installed AlphaFold v2.2 from Github (https://github.com/deepmind/alphafold) in May 2022 and v.2.3 in February 2023. Both versions of AlphaFold were installed on a local computing cluster. During the structure prediction or feature preparation step in the AlphaFold pipeline, 15 cases were unable to complete because of GPU and memory limitations out of a total of 444 test cases.

For generating unbound antibody and antigen structures, we employed AlphaFold in Multimer setting when the input consisted of a heavy-light chain antibody or a multimeric antigen. Alternatively, the Monomer setting was utilized when the input was a single chain. A template date cutoff of 2018-04-30 was applied to avoid template overlap with benchmarking set.

To generate AlphaFold predictions without the use of MSAs (corresponding to single-sequence modeling), we modified “all_seq_msa_features” variable of chain features, to include only the query sequence. To use custom templates, we adapted the template featurization function from Motmaen et al.^44^ (https://github.com/phbradley/alphafold_finetune/blob/main/predict_utils.py).

AlphaFold modeling in ColabFold^27^ was performed with ColabFold version 1.3.0 (commit 26de12d3afb5f85d49d0c7db1b9371f034388395), installed on a local computing cluster using scripts from Github (https://github.com/YoshitakaMo/localcolabfold). During ColabFold AlphaFold modeling, MSA was built by querying the MMseqs2 MSA server using unpaired and paired MSA. To generate a total of 25 predictions per complex, modifications were made to “load_models_and_params” function, utilizing a different random seed for each prediction, producing five predictions per AlphaFold model parameter.

Unless otherwise specified, a template date cutoff of April 30, 2018 was applied for benchmarking AlphaFold v.2.2 and ColabFold, and a template date cutoff of September 30, 2021 was applied for benchmarking of AlphaFold v.2.3, to avoid using bound structures as template.

AlphaFold and ColabFold modeling runs were performed using NVIDIA Titan RTX and Quadro 6000 GPUs.

### Complex model accuracy assessment

We assessed antibody-antigen complex model accuracy using DockQ^36^, which was downloaded from GitHub (https://github.com/bjornwallner/DockQ). Antibody-antigen complex model accuracy was computed by DockQ using the experimentally determined antibody-antigen complex structures obtained from the PDB. DockQ calculates interface backbone RMSD (I-RMSD), ligand backbone RMSD (L-RMSD), fraction of native contacts (fnat), DockQ score, as well as the CAPRI (Critical Assessment of PRediction of Interactions) accuracy level, which assigns the model into one of four discrete accuracy classes: Incorrect, Acceptable, Medium, and High, based on the model’s similarity to the native structure.^26^

### CDR loop accuracy analysis

The complementarity determining regions (CDRs) and the framework regions of antibodies were identified by AHo numbering^53^, assigned using ANARCI software^54^. The CDR loops were defined as: residues 24-42 (CDR1), 57-76 (CDR2), and 107-138 (CDR3).

ProFit v 3.1^55^ was used to calculate backbone RMSDs between modeled and experimentally determined CDR loop structures, after superposing the modeled antibody structures onto the experimentally resolved structures by the framework residues.

### MSA depth calculation

The MSA of the antibody-antigen complex was retrieved from the ’msa’ key value in the feature dictionary (“feature_dict” variable). Given that the MSA values of the ‘msa’ key are encoded in AlphaFold residue IDs, we converted the amino acids back to the one-letter amino acid type using “ID_TO_HHBLITS_AA” dictionary, and replaced gaps (denoted by “-“) by “U” for downstream MSA depth calculation. Number of effective sequences (N_eff_) was calculated by CD-Hit^56^. For consistency with the AlphaFold MSA N_eff_ calculation scheme^22^, we used an identity cutoff of 80% in CD-HIT to calculate nonredundant sequence clusters. MSA depth was successfully calculated for 428 out of 429 antibody-antigen complexes, with one failure due to a technical issue.

### Interface glycans

Glycans were identified through inspection of HETATM records in experimentally resolved structure coordinates. Biological assembly structures (defined by the PDB entry) of the antibody-antigen complexes were inspected, and cases with hetero-atoms matching the “saccharide” classification identified from PDB ligands summary pages (from wwPDB’s Chemical Component Dictionary) within 4.5 Å of the antibody were identified as positive hits.

### Figures and statistical analysis

PyMOL version 2.4 (Schrodinger, Inc.) was used to generate figures of antibody-antigen complex structures. The ggplot2^57^ package in R (r-project.org) was utilized to generate box plots, line plots, and bar plots. Pearson correlations and their corresponding p values were calculated using the ggpubr package in R, while the Wilcoxon rank-sum test was performed using the ggsignif package in R. Binary and multi-class ROC curves with AUC values were calculated using the pROC^58^ and multiROC^59^ packages in R, respectively.

### Interface pLDDT calculation

To determine the interface pLDDT (I-pLDDT), we computed the average pLDDT value for all residues at the antibody-antigen interface. Interface residues were defined as any residue with a non-hydrogen atom within 4.0 Å of the binding partner. An I-pLDDT score of 30 was assigned to predictions with no antibody-antigen interface residues.

### Antibody-antigen complex scoring and native complex relaxation

The “InterfaceAnalyzer” executable in Rosetta^32^ (v.3.12) was employed to calculate interface energetic scores, using the Rosetta Energy Function 2015 (REF15) scoring function^60^ and default parameters. Prior to scoring, structural relaxation was performed on native antibody-antigen complex obtained from PDB using the FastRelax protocol^61^ (“relax” executable) in Rosetta using the following flags:

-ignore_unrecognized_res

-relax:constrain_relax_to_start_coords

-relax:coord_constrain_sidechains

-relax:ramp_constraints false

-ex1

-ex2

-use_input_sc

-no_optH false

-flip_HNQ

-overwrite

-nstruct 1

### Rigid-body docking model generation with ZDOCK and IRAD rescoring

To establish a rigid-body docking baseline, ZDOCK version 3.0.2^28^ was used to generate antibody-antigen docking models using unbound or bound input structures. Unbound antibody and antigen input structures for the ZDOCK algorithm were generated by AlphaFold, using the above-mentioned protocol. Bound antibody and antigen input structures were extracted from experimentally resolved antibody-antigen complex structures downloaded from PDB, with HETATMs removed. Through dense rotational sampling (“-D” flag in ZDOCK), a total of 54,000 predictions per complex were generated. Subsequently, the docking poses were scored and reranked using the IRAD scoring function^29^. Finally, the top predictions were scored with DockQ to determine the modeling accuracy.

### TM-score calculation

To provide an assessment of antigen prediction accuracy, we employed the TM-score executable^33^ to calculate the TM-score, comparing the structural similarity between the antigen chain(s) in the experimentally resolved antibody-antigen complex structure, and the antigen prediction in the antibody-antigen complex predictions. Prior to running TM-score on the antigen chains, residues that were not experimentally resolved, or absent from the experimentally resolved structure, were removed from the antigen prediction.

## Data Availability

Modified AlphaFold code and analysis scripts are available on Github: https://github.com/piercelab/alphafold_v2.2_customize. AlphaFold2.2, AlphaFold2.3, and ColabFold antibody-antigen models generated in this study are available for download at: https://piercelab.ibbr.umd.edu/af_abag_benchmarking.html.

## Acknowledgements

We are grateful to the information technology team at the Institute for Bioscience and Biotechnology Research, including Gale Lane who provided valuable assistance with high performance computing resources. Members of the Pierce lab, including Helder Veras Ribeiro-Filho, provided insightful discussions and suggestions. We also express our gratitude to John Moult, Brandon Feng, Mike Song, Sergey Ovchinnikov (Harvard University), and John Jumper (Deepmind) for helpful discussions. This work was supported by National Institutes of Health grant R35 GM144083 to B.G.P.

## Author Contributions

R.Y. and B.G.P. conceived the study. R.Y. performed modeling simulations, analyzed results, and drafted the manuscript. R.Y. and B.G.P. edited the manuscript.

## References

1 Chothia, C. & Lesk, A. M. Canonical structures for the hypervariable regions of immunoglobulins. J Mol Biol 196, 901–917 (1987). https://doi.org:10.1016/0022-2836(87)90412-8

2 Sela-Culang, I., Kunik, V. & Ofran, Y. The structural basis of antibody-antigen recognition. Front Immunol 4, 302 (2013). https://doi.org:10.3389/fimmu.2013.00302

3 Nelson, A. L., Dhimolea, E. & Reichert, J. M. Development trends for human monoclonal antibody therapeutics. Nat Rev Drug Discov 9, 767–774 (2010). https://doi.org:10.1038/nrd3229

4 Carter, P. J. Potent antibody therapeutics by design. Nat Rev Immunol 6, 343–357 (2006). https://doi.org:10.1038/nri1837

5 Scott, A. M., Wolchok, J. D. & Old, L. J. Antibody therapy of cancer. Nat Rev Cancer 12, 278–287 (2012). https://doi.org:10.1038/nrc3236

6 Rappuoli, R., Bottomley, M. J., D’Oro, U., Finco, O. & De Gregorio, E. Reverse vaccinology 2.0: Human immunology instructs vaccine antigen design. J Exp Med 213, 469–481 (2016). https://doi.org:10.1084/jem.20151960

7 Li, Y., Li, H., Yang, F., Smith-Gill, S. J. & Mariuzza, R. A. X-ray snapshots of the maturation of an antibody response to a protein antigen. Nat Struct Biol 10, 482–488 (2003).

8 Barnes, C. O. et al. SARS-CoV-2 neutralizing antibody structures inform therapeutic strategies. Nature 588, 682–687 (2020). https://doi.org:10.1038/s41586-020-2852-1

9 Zhou, T. et al. Structural Repertoire of HIV-1-Neutralizing Antibodies Targeting the CD4 Supersite in 14 Donors. Cell 161, 1280–1292 (2015). https://doi.org:10.1016/j.cell.2015.05.007

10 Dreyfus, C. et al. Highly conserved protective epitopes on influenza B viruses. Science 337, 1343–1348 (2012). https://doi.org:10.1126/science.1222908

11 Haidar, J. N. et al. A universal combinatorial design of antibody framework to graft distinct CDR sequences: a bioinformatics approach. Proteins 80, 896–912 (2012). https://doi.org:10.1002/prot.23246

12 Hanf, K. J. et al. Antibody humanization by redesign of complementarity-determining region residues proximate to the acceptor framework. Methods 65, 68–76 (2014). https://doi.org:10.1016/j.ymeth.2013.06.024

13 Graham, B. S., Gilman, M. S. A. & McLellan, J. S. Structure-Based Vaccine Antigen Design. Annu Rev Med 70, 91–104 (2019). https://doi.org:10.1146/annurev-med-121217-094234

14 Georgiou, G. et al. The promise and challenge of high-throughput sequencing of the antibody repertoire. Nat Biotechnol 32, 158–168 (2014). https://doi.org:10.1038/nbt.2782

15 Li, Z., Woo, C. J., Iglesias-Ussel, M. D., Ronai, D. & Scharff, M. D. The generation of antibody diversity through somatic hypermutation and class switch recombination. Genes Dev 18, 1–11 (2004). https://doi.org:10.1101/gad.1161904

16 Vreven, T. et al. Updates to the Integrated Protein-Protein Interaction Benchmarks: Docking Benchmark Version 5 and Affinity Benchmark Version 2. J Mol Biol 427, 3031–3041 (2015). https://doi.org:10.1016/j.jmb.2015.07.016

17 Sircar, A. & Gray, J. J. SnugDock: paratope structural optimization during antibody-antigen docking compensates for errors in antibody homology models. PLoS computational biology 6, e1000644 (2010). https://doi.org:10.1371/journal.pcbi.1000644

18 Brenke, R. et al. Application of asymmetric statistical potentials to antibody-protein docking. Bioinformatics 28, 2608–2614 (2012). https://doi.org:10.1093/bioinformatics/bts493

19 Krawczyk, K., Baker, T., Shi, J. & Deane, C. M. Antibody i-Patch prediction of the antibody binding site improves rigid local antibody-antigen docking. Protein Eng Des Sel 26, 621–629 (2013). https://doi.org:10.1093/protein/gzt043

20 Ambrosetti, F., Jimenez-Garcia, B., Roel-Touris, J. & Bonvin, A. Modeling Antibody-Antigen Complexes by Information-Driven Docking. Structure 28, 119–129 e112 (2020). https://doi.org:10.1016/j.str.2019.10.011

21 Guest, J. D. et al. An expanded benchmark for antibody-antigen docking and affinity prediction reveals insights into antibody recognition determinants. Structure 29, 606–621 e605 (2021). https://doi.org:10.1016/j.str.2021.01.005

22 Jumper, J. et al. Highly accurate protein structure prediction with AlphaFold. Nature 596, 583–589 (2021). https://doi.org:10.1038/s41586-021-03819-2

23 Jumper, J. et al. Applying and improving AlphaFold at CASP14. Proteins (2021). https://doi.org:10.1002/prot.26257

24 Evans, R., et al. Protein complex prediction with AlphaFold-Multimer. bioRxiv (2021).

25 Yin, R., Feng, B. Y., Varshney, A. & Pierce, B. G. Benchmarking AlphaFold for protein complex modeling reveals accuracy determinants. Protein Sci 31, e4379 (2022). https://doi.org:10.1002/pro.4379

26 Lensink, M. F., Nadzirin, N., Velankar, S. & Wodak, S. J. Modeling protein-protein, protein-peptide, and protein-oligosaccharide complexes: CAPRI 7th edition. Proteins 88, 916–938 (2020). https://doi.org:10.1002/prot.25870

27 Mirdita, M. et al. ColabFold: making protein folding accessible to all. Nat Methods 19, 679–682 (2022). https://doi.org:10.1038/s41592-022-01488-1

28 Pierce, B. G., Hourai, Y. & Weng, Z. Accelerating protein docking in ZDOCK using an advanced 3D convolution library. PLoS One 6, e24657 (2011). https://doi.org:10.1371/journal.pone.0024657

29 Vreven, T., Hwang, H. & Weng, Z. Integrating atom-based and residue-based scoring functions for protein-protein docking. Protein Sci 20, 1576–1586 (2011). https://doi.org:10.1002/pro.687

30 Guest, J. D. et al. An expanded benchmark for antibody-antigen docking and affinity prediction reveals insights into antibody recognition determinants. Structure (2021). https://doi.org:10.1016/j.str.2021.01.005

31 Kappler, K. & Hennet, T. Emergence and significance of carbohydrate-specific antibodies. Genes Immun 21, 224–239 (2020). https://doi.org:10.1038/s41435-020-0105-9

32 Leman, J. K. et al. Macromolecular modeling and design in Rosetta: recent methods and frameworks. Nat Methods 17, 665–680 (2020). https://doi.org:10.1038/s41592-020-0848-2

33 Zhang, Y. & Skolnick, J. Scoring function for automated assessment of protein structure template quality. Proteins 57, 702–710 (2004). https://doi.org:10.1002/prot.20264

34 Xu, J. & Zhang, Y. How significant is a protein structure similarity with TM-score = 0.5? Bioinformatics 26, 889–895 (2010). https://doi.org:10.1093/bioinformatics/btq066

35 Bryant, P., Pozzati, G. & Elofsson, A. Improved prediction of protein-protein interactions using AlphaFold2. Nat Commun 13, 1265 (2022). https://doi.org:10.1038/s41467-022-28865-w

36 Johansson-Akhe, I. & Wallner, B. Improving peptide-protein docking with AlphaFold-Multimer using forced sampling. Front Bioinform 2, 959160 (2022). https://doi.org:10.3389/fbinf.2022.959160

37 DeepMind. AlphaFold v2.3.0, <https://github.com/deepmind/alphafold/blob/main/docs/technical_note_v2.3.0.md> (2022).

38 Yin, R., Feng, B. Y., Varshney, A. & Pierce, B. G. Benchmarking AlphaFold for protein complex modeling reveals accuracy determinants. bioRxiv, 2021.2010.2023.465575 (2021). https://doi.org:10.1101/2021.10.23.465575

39 Deepmind. AlphaFold v2.3.0 technical note, <https://github.com/deepmind/alphafold/blob/main/docs/technical_note_v2.3.0.md> (2022).

40 Björn, W. AFsample: Improving Multimer Prediction with AlphaFold using Aggressive Sampling. bioRxiv, 2022.2012.2020.521205 (2023). https://doi.org:10.1101/2022.12.20.521205

41 Sala, D., Engelberger, F., McHaourab, H. S. & Meiler, J. Modeling conformational states of proteins with AlphaFold. Curr Opin Struct Biol 81, 102645 (2023). https://doi.org:10.1016/j.sbi.2023.102645

42 Ziyao, L. et al. Uni-Fold: An Open-Source Platform for Developing Protein Folding Models beyond AlphaFold. bioRxiv, 2022.2008.2004.502811 (2022). https://doi.org:10.1101/2022.08.04.502811

43 Gustaf, A. et al. OpenFold: Retraining AlphaFold2 yields new insights into its learning mechanisms and capacity for generalization. bioRxiv, 2022.2011.2020.517210 (2022). https://doi.org:10.1101/2022.11.20.517210

44 Motmaen, A. et al. Peptide-binding specificity prediction using fine-tuned protein structure prediction networks. Proc Natl Acad Sci U S A 120, e2216697120 (2023). https://doi.org:10.1073/pnas.2216697120

45 Zheng, W. et al. Protein structure prediction using deep learning distance and hydrogen-bonding restraints in CASP14. Proteins 89, 1734–1751 (2021). https://doi.org:10.1002/prot.26193

46 Lensink, M., et al. (Authorea Preprints, 2023).

47 Bo, C. et al. Improved the Protein Complex Prediction with Protein Language Models. bioRxiv, 2022.2009.2015.508065 (2022). https://doi.org:10.1101/2022.09.15.508065

48 Ruffolo, J. A., Gray, J. J. & Sulam, J. Deciphering antibody affinity maturation with language models and weakly supervised learning. arXiv preprint arXiv:2112.07782 (2021).

49 Abanades, B. et al. ImmuneBuilder: Deep-Learning models for predicting the structures of immune proteins. Commun Biol 6, 575 (2023). https://doi.org:10.1038/s42003-023-04927-7

50 Ruffolo, J. A., Chu, L. S., Mahajan, S. P. & Gray, J. J. Fast, accurate antibody structure prediction from deep learning on massive set of natural antibodies. Nat Commun 14, 2389 (2023). https://doi.org:10.1038/s41467-023-38063-x

51 Camacho, C. et al. BLAST+: architecture and applications. BMC bioinformatics 10, 421 (2009). https://doi.org:10.1186/1471-2105-10-421

52 Zhu, J. & Weng, Z. FAST: a novel protein structure alignment algorithm. Proteins 58, 618–627 (2005). https://doi.org:10.1002/prot.20331

53 Honegger, A. & Pluckthun, A. Yet another numbering scheme for immunoglobulin variable domains: an automatic modeling and analysis tool. J Mol Biol 309, 657–670 (2001). https://doi.org:10.1006/jmbi.2001.4662

54 Dunbar, J. & Deane, C. M. ANARCI: antigen receptor numbering and receptor classification. Bioinformatics 32, 298–300 (2016). https://doi.org:10.1093/bioinformatics/btv552

55 Martin, A. C. & Porter, C. T. ProFit Version 3.1, <http://www.bioinf.org.uk/software/profit/> (2009).

56 Fu, L., Niu, B., Zhu, Z., Wu, S. & Li, W. CD-HIT: accelerated for clustering the next-generation sequencing data. Bioinformatics 28, 3150-3152 (2012). https://doi.org:10.1093/bioinformatics/bts565

57 Wickham, H. ggplot2: Elegant Graphics for Data Analysis. (Springer-Verlag New York, 2016).

58 Robin, X. et al. pROC: an open-source package for R and S+ to analyze and compare ROC curves. BMC bioinformatics 12, 77 (2011). https://doi.org:10.1186/1471-2105-12-77

59 Wei, R. & Wang, J. multiROC: Calculating and Visualizing ROC and PR Curves Across Multi-Class assifications. (2018).

60 Alford, R. F. et al. The Rosetta All-Atom Energy Function for Macromolecular Modeling and Design. J Chem Theory Comput 13, 3031–3048 (2017). https://doi.org:10.1021/acs.jctc.7b00125

61 Conway, P., Tyka, M. D., DiMaio, F., Konerding, D. E. & Baker, D. Relaxation of backbone bond geometry improves protein energy landscape modeling. Protein Sci 23, 47–55 (2014). https://doi.org:10.1002/pro.2389

62 Stranges, P. B. & Kuhlman, B. A comparison of successful and failed protein interface designs highlights the challenges of designing buried hydrogen bonds. Protein Sci 22, 74–82 (2013). https://doi.org:10.1002/pro.2187

