## Supplementary material for "Evaluation of AlphaFold Antibody-Antigen Modeling with Implications for Improving Predictive Accuracy": Figures S1-S14, Table S1

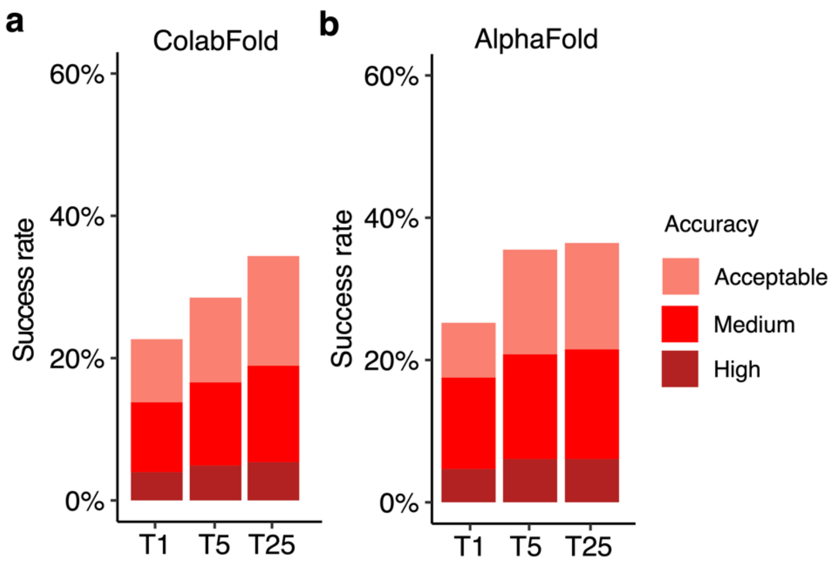


**Figure S1. Antibody-antigen modeling success by ColabFold and AlphaFold.** Antibody-antigen modeling success comparison of **a** ColabFold and **b** AlphaFold on 428 antibody-antigen complexes for which both algorithms successfully generated predictions. Templates released on or before April 30, 2018, were allowed during modeling. For each complex, 25 predictions were generated, and were ranked by AlphaFold model confidence score. Antibody-antigen predictions were evaluated for complex modeling accuracy using CAPRI criteria for High, Medium and Acceptable accuracy. The success rate was calculated based on the percentage of cases that had at least one model among their top N predictions that met a specified level of CAPRI accuracy. Bars were colored as per CAPRI accuracy criteria.

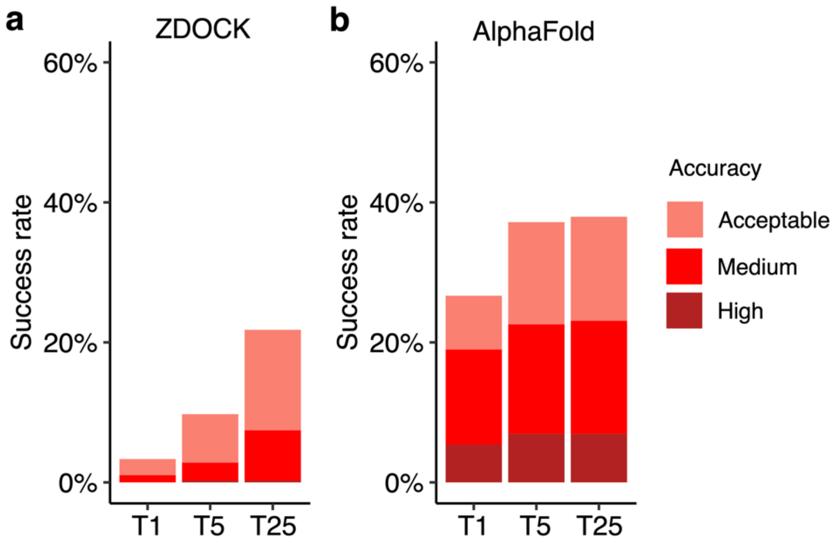


**Figure S2. Antibody-antigen modeling success by ZDOCK and AlphaFold.** Antibody-antigen modeling success comparison of **a** ZDOCK (version 3.0.2)[6] **b** AlphaFold on 390 antibody-antigen complexes. AlphaFold was used to generate unbound antibody and antigen structure inputs for ZDOCK. Templates released on or before April 30, 2018, were allowed during modeling. Only the top-ranked prediction, based on AlphaFold's model confidence score, was used as input. ZDOCK docking was performed only on inputs with a minimum antibody or antigen pLDDT score above 80 to ensure input model quality. ZDOCK dense sampling was employed, generating in 54,000 predictions per complex, which were subsequently ranked using IRAD[7] scores. Antibody-antigen predictions were evaluated for complex modeling accuracy using CAPRI criteria for High, Medium and Acceptable accuracy. For each complex, 25 predictions were generated, and were ranked by AlphaFold model confidence score. The success rate was calculated based on the percentage of cases that had at least one model among their top N predictions that met a specified level of CAPRI accuracy. Bars were colored as per CAPRI accuracy criteria.

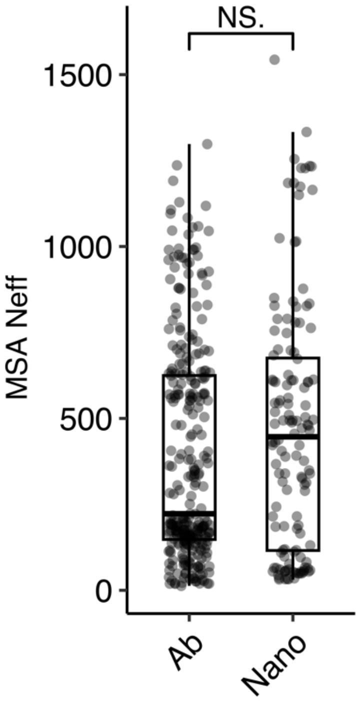


**Figure S3. Distribution of MSA depth (Neff) grouped by antibody type**. Based on the antibody type, complexes are categorized into heavy-light chain antibody-antigen complexes (Ab, N=297), or nanobody/VHH-antibody complexes (Nano, N=132). Statistical significance values (Wilcoxon rank-sum test) were calculated between MSA depth for antibody targets versus nanobody targets, as noted at top (NS.: not significant, $p > 0.05$).

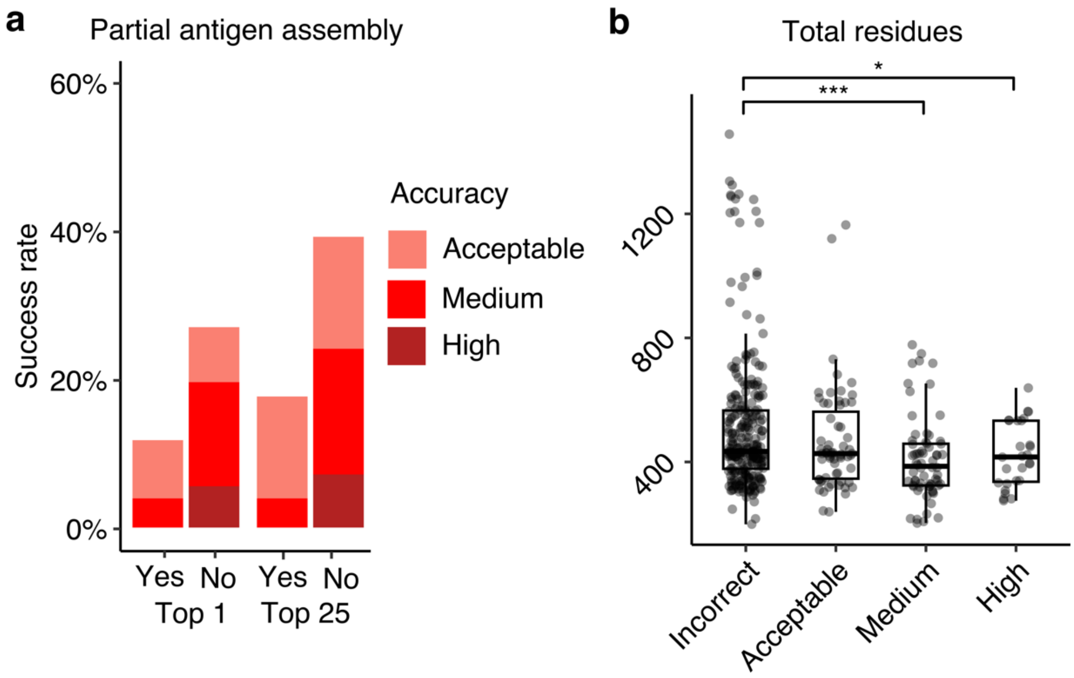


**Figure S4. Antibody-antigen modeling success determinants. a** Partial versus full antigen assembly input. Complexes were classified as either "Yes" (N=51) or "No" (N=378) to indicate whether a partial antigen assembly was modeled, meaning that the antigen was modeled without additional chains that are present in the full PDB bioassembly. T1 and T25 denote AlphaFold modeling accuracy in top 1 (ranked by AlphaFold model confidence score) and in all 25 predictions of the complex. Bars are colored by CAPRI criteria. **b** Total number of residues in the complex grouped by AlphaFold modeling success. The modeling success is defined as the highest CAPRI criteria prediction in the complex, considering all 25 predictions. Statistical significance values (Wilcoxon rank-sum test) were calculated between total residue counts for sets of cases with Incorrect versus Medium and Incorrect versus High CAPRI accuracy predictions, as noted at top (*$p \leq 0.05$, ***$p \leq 0.001$).

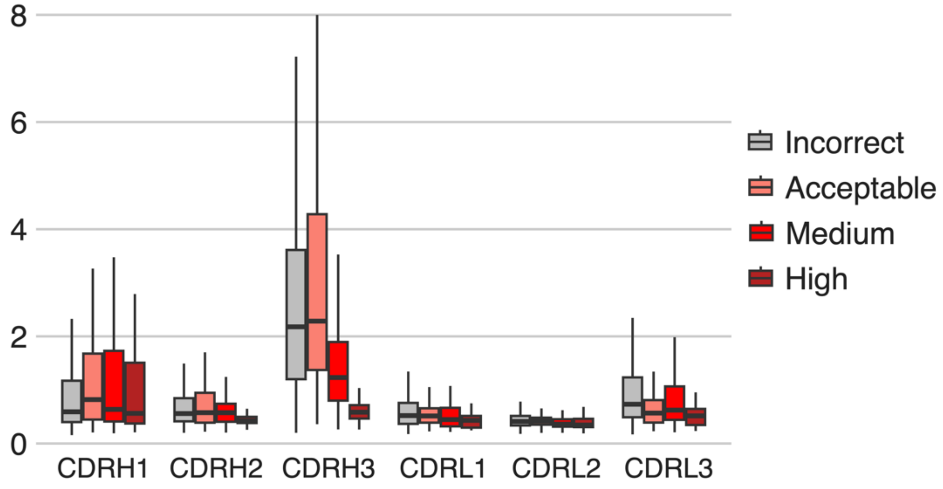


**Figure S5. Distribution of CDR modeling accuracy, grouped by CDR type and by AlphaFold modeling accuracy.** AlphaFold modeling accuracy is defined as the accuracy of highest CAPRI criteria prediction in the complex, considering all 25 predictions. For CDRH1, CDRH2 and CDRH3, the numbers of data points in each category are 263, 60, 62, 25 for Incorrect, Acceptable, Medium and High success groups. For CDRL1, CDRL2 and CDRL3, the numbers of data points in each category are 194, 45, 39, 10 for Incorrect, Acceptable, Medium and High success groups. Bars colored by CAPRI criteria.

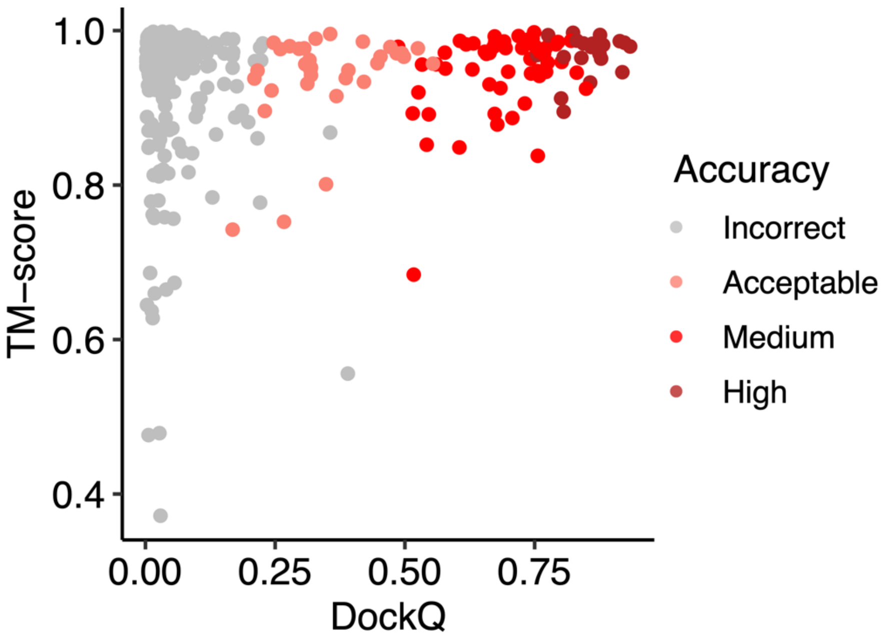


**Figure S6. Relationship between antigen modeling accuracy and complex prediction accuracy.** Top-ranked predictions of 429 complexes generated by AlphaFold are represented as data points. The antigen accuracy is measured by TM-score of the top-ranked prediction, and the complex model accuracy measured by DockQ score. If the antigen has multiple chains, the minimum TM-score of all antigen chains is selected as the antigen TM-score. Data points are colored by the CAPRI criteria.

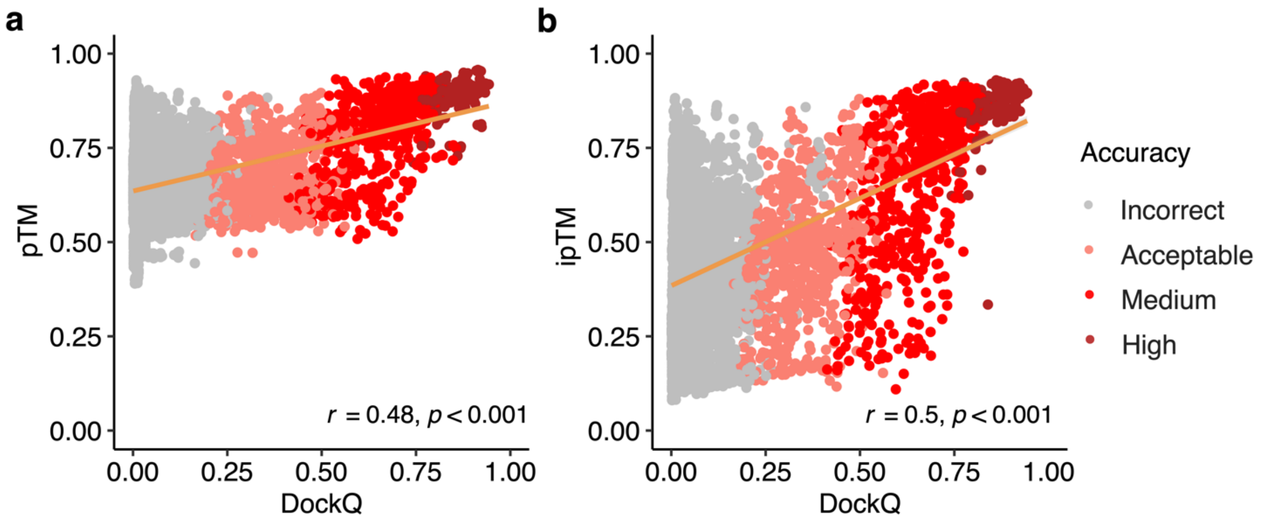


**Figure S7**. **Relationship between model confidence scores and model accuracy.** Scatter plots depicting the association between the **a** pTM, **b** ipTM scores and the DockQ scores. In the scatter plots, all 25 models representing 429 complexes are depicted as data points, with their colors indicating the model quality according to CAPRI criteria. The orange line represents the linear regression, and the lower right corner of the scatter plots displays the Pearson's correlation coefficients and correlation p-values.

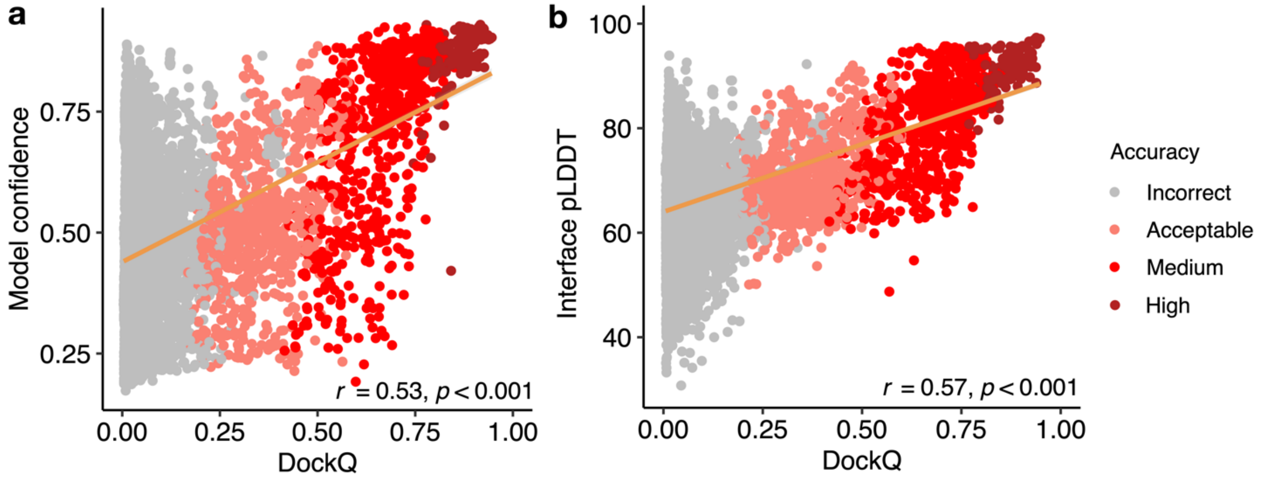


**Figure S8**. **Relationship between model scores and model accuracy.** Scatter plots depicting the association between the **a** model confidence, **b** interface pLDDT scores and the DockQ scores. A total of 10,344 data points were present in each scatter plot, which include all 25 models representing 429 complexes, excluding models without side-chain contacts within 4 Å across the antibody-antigen interface. Data points are colors indicating the model quality according to CAPRI criteria. The orange line represents the linear regression, and the lower right corner of the scatter plots displays the Pearson's correlation coefficients and correlation p-values.

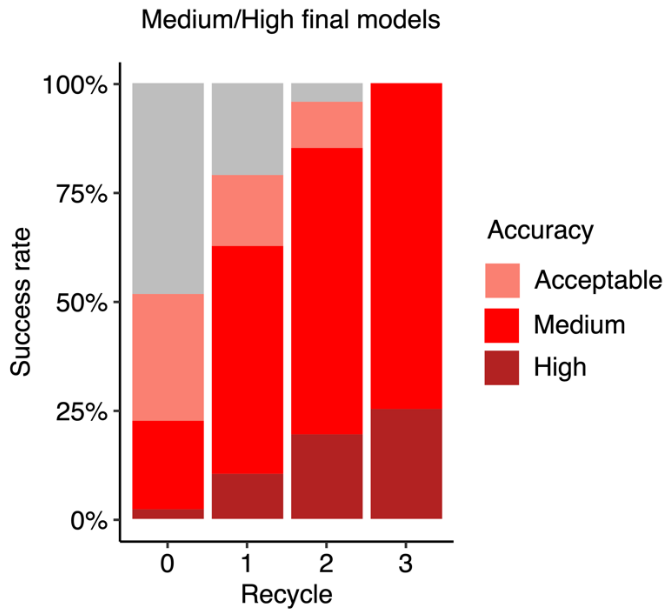


**Figure S9. Analysis of antibody-antigen predictions accuracy across recycling iterations.** The modeling success of complexes at each recycle focusing on a subset of predictions that reached Medium or higher accuracy after 3 recycling iterations (N=109). Recycle=0 denotes the state of the prediction before recycling iterations begin.

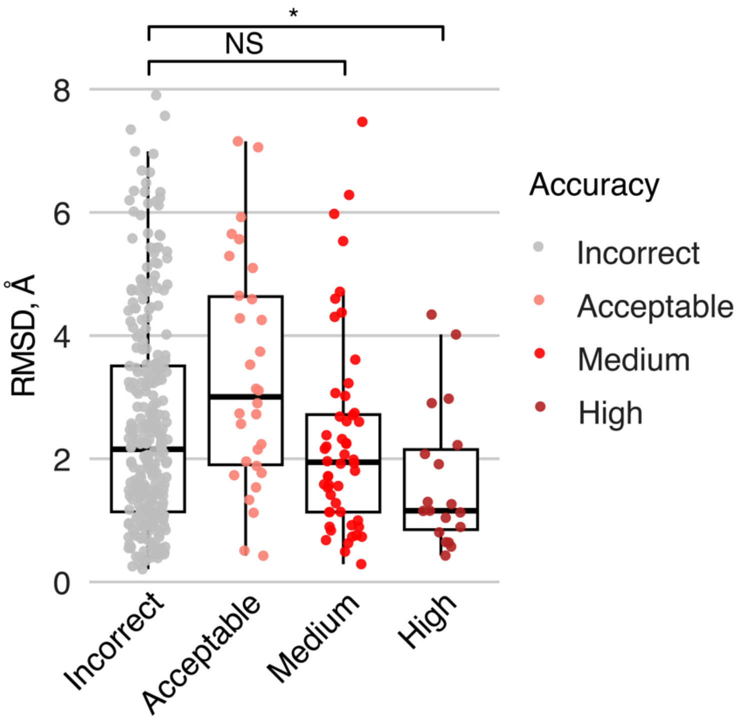


**Figure S10. The distribution of CDRH3 modeling accuracy in top-ranked unbound antibody model grouped by top-ranked complex modeling success.** The modeling success is CAPRI criteria of top-ranked complex prediction generated by AlphaFold. The CDRH3 RMSD measures the RMSD of the top-ranked unbound antibody prediction generated by AlphaFold. Numbers of data points in Incorrect, Acceptable, Medium and High categories are 306, 31, 52, 19. Statistical significance values (Wilcoxon rank-sum test) were calculated between RMSD values for sets of cases with Incorrect versus Medium and Incorrect versus High CAPRI accuracy predictions, as noted at top (NS: $p > 0.05$, *$p \leq 0.05$).

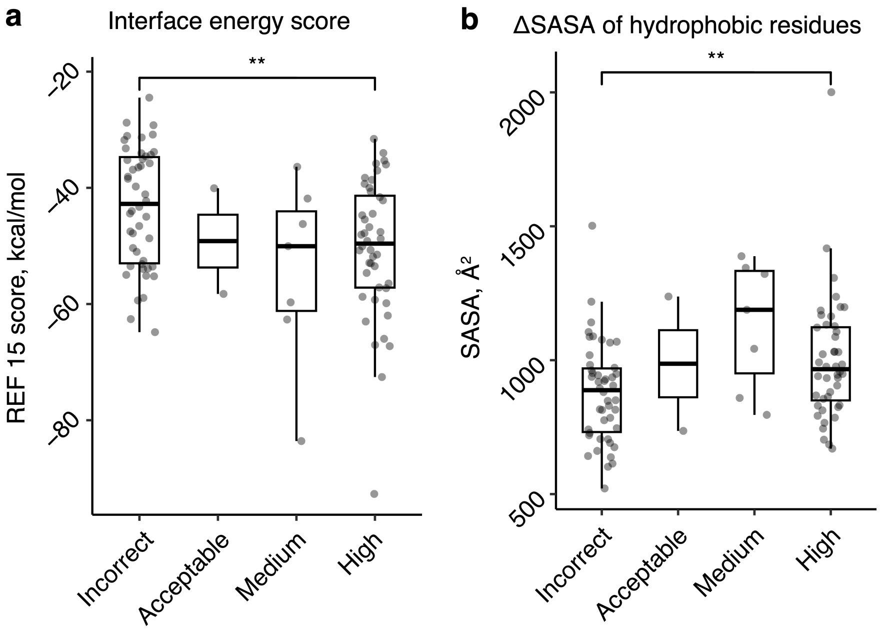


**Figure S11. Antibody-antigen modeling success determinants of AlphaFold with bound antibody and antigen structures as templates.** Distribution of **a** interface energy score and **b** Δ solvent-accessible surface area (ΔSASA) of hydrophobic part of the antibody-antigen interface by complex modeling success by AlphaFold when using bound antibody and antigen structures as templates. The modeling success is defined as the highest CAPRI criteria prediction in the complex, considering all 5 predictions. Numbers of data points in Incorrect, Acceptable, Medium and High categories are 46, 2, 7 and 45. Statistical significance values (Wilcoxon rank-sum test) were calculated between scores for sets of cases with Incorrect versus High CAPRI accuracy predictions, as noted at top (**$p \leq 0.01$).

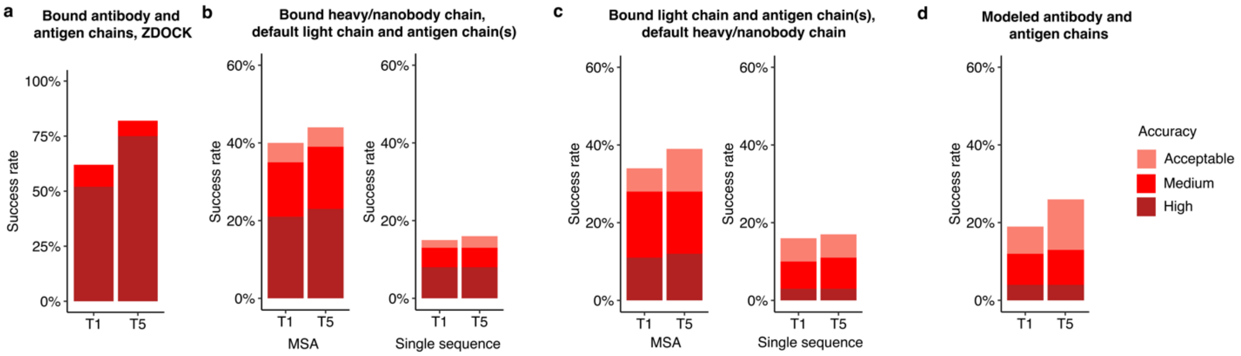


**Figure S12. Antibody-antigen modeling success using varying template inputs.** Antibody-antigen modeling success of **a** ZDOCK (version 3.0.2) with IRAD re-ranking of dense-sampling predictions (54,000 predictions per complex), utilizing bound antibody and bound antigen chains as docking input, and of AlphaFold by utilizing **b** bound heavy/nanobody chain, and default light chain and antigen chains as templates, **c** bound light chain, and default heavy/nanobody chain and antigen chains as templates, **d** antibody and antigen chains modeled by AlphaFold as templates. Benchmarking was performed on a total of 100 antibody-antigen complexes. The success rate was calculated based on the percentage of cases that had at least one model among their top N predictions that met a specified level of CAPRI accuracy. Bars were colored by CAPRI accuracy criteria.

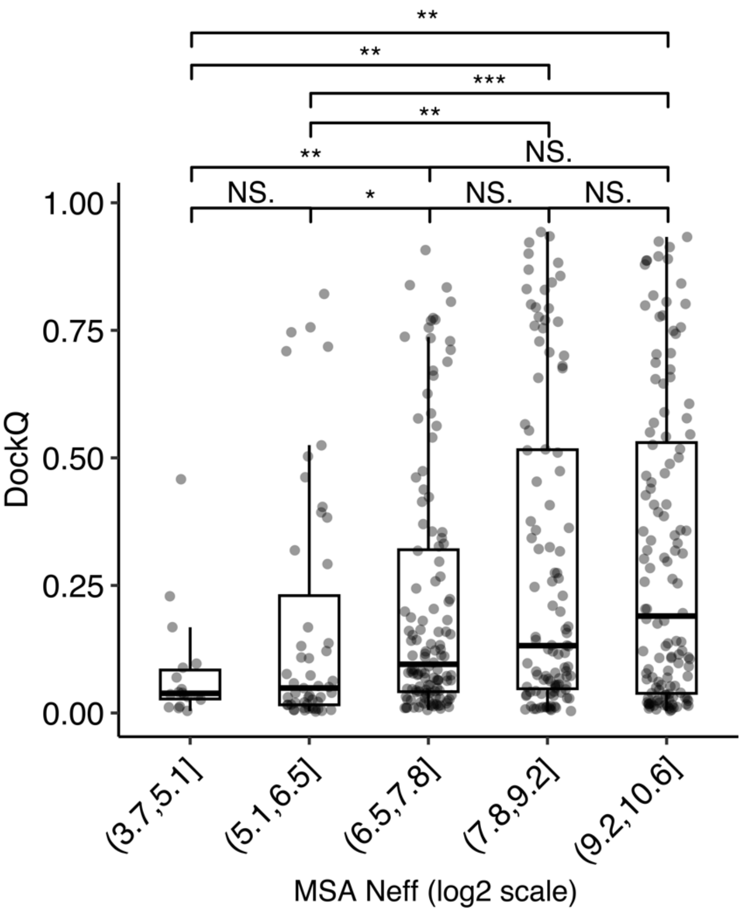


**Figure S13. Distribution of DockQ scores grouped by ranges of AlphaFold MSA depth (log2 scale).** The DockQ scores selected for each individual data points are the highest DockQ score of all 25 complex predictions generated by AlphaFold. Numbers of data points in (3.69,5.08], (5.08,6.46], (6.46,7.84], (7.84,9.21] and (9.21,10.6] Neff ranges are 18, 15, 124, 107, 128 respectively. Statistical significance values (Wilcoxon rank-sum test) were calculated between DockQ scores for sets of cases with varying ranges of MSA depth, as noted at top (NS.: $p > 0.05$, $^*p \leq 0.005$, $^{**}p \leq 0 .001$, $^{***}p \leq 0.001$).

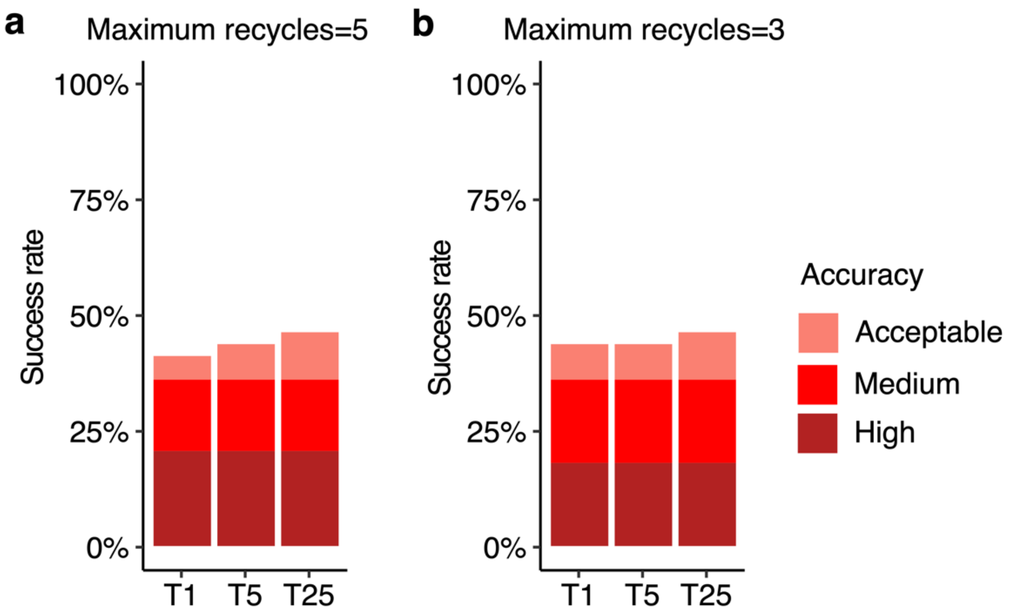


**Figure S14. Antibody-antigen modeling success comparison of AlphaFold (v.2.3) using a varying number of maximum recycles.** The maximum number of recycling iterations were set to **a** 5 and **b** 3, and tested on 39 antibody-antigen complexes. Templates released on or before September 30, 2021, were allowed during modeling. For each complex, 25 predictions were generated, and were ranked by AlphaFold model confidence score. Antibody-antigen predictions were evaluated for complex modeling accuracy using CAPRI criteria for High, Medium and Acceptable accuracy. The success rate was calculated based on the percentage of cases that had at least one model among their top N predictions that met a specified level of CAPRI accuracy. Bars were colored as per CAPRI accuracy criteria.

**Table S1. Antibody-antigen structures used for AlphaFold benchmarking.**

| PDB[1] | Heavy chain | Light chain | Antigen chain(s) | Release Date | v.2.3.0 set[2] | 100 subset |
|---|---|---|---|---|---|---|
| 6was | H | L | J | 3/31/21 | | yes |
| 6p50 | H | L | C | 9/4/19 | | |
| 6urm | H | L | F | 9/16/20 | | |
| 6umg | h | l | cr | 2/12/20 | | |
| 6a0z | H | L | A | 6/20/18 | | yes |
| 7daa | H | L | A | 10/20/21 | yes | yes |
| 6s8j | P | O | C | 2/12/20 | | |
| 6s8i | P | O | C | 2/12/20 | | |
| 7lf7 | A | B | M | 8/4/21 | | |
| 6urh | H | L | C | 3/18/20 | | |
| 7lfb | H | L | X | 8/4/21 | | |
| 6ktr | A | B | C | 7/8/20 | | |
| 6meh | H | L | C | 11/21/18 | | |
| 6vel | H | L | C | 1/29/20 | | yes |
| 6y9b | I | M | A | 5/20/20 | | |
| 6xkq | H | L | A | 10/14/20 | | |
| 6o9h | H | L | D | 1/22/20 | | |
| 6yio | H | L | B | 11/11/20 | | |
| 7l7r | B | A | G | 12/1/21 | yes | |
| 7l7r | D | C | G | 12/1/21 | yes | yes |
| 6svl | A | B | C | 11/27/19 | | |
| 6okm | H | L | R | 8/28/19 | | |
| 6z3p | H | L | CAB | 9/2/20 | | |
| 6u36 | H | L | B | 11/6/19 | | |
| 6jbt | H | L | F | 6/19/19 | | |
| 6wbv | H | L | A | 9/9/20 | | |
| 7jmp | H | L | A | 8/26/20 | | yes |
| 6wo5 | G | I | F | 8/19/20 | | yes |
| 7lfa | B | D | A | 8/4/21 | | |
| 7np1 | H | L | A | 11/17/21 | | |
| 6hig | H | L | B | 6/5/19 | | |
| 6vyh | D | C | A | 11/11/20 | | |
| 6k65 | H | L | A | 8/14/19 | | |
| 6umx | h | l | B | 2/26/20 | | |
| 7nx3 | B | C | F | 10/27/21 | yes | |
| 7jv6 | C | D | B | 10/14/20 | | |
| 6xqw | H | L | E | 3/3/21 | | |

| | | | | | |
|---|---|---|---|---|---|
| 7jtg | A | B | E | 3/10/21 | yes |
| 6o9i | D | E | C | 1/22/20 | yes |
| 7e7x | H | L | A | 6/9/21 | |
| 6wo5 | A | B | E | 8/19/20 | |
| 6mvl | H | L | A | 10/23/19 | |
| 7kqg | B | C | A | 12/16/20 | |
| 6glw | H | L | A | 6/5/19 | |
| 6gku | H | L | A | 6/5/19 | |
| 7lue | H | L | A | 6/16/21 | |
| 7k9j | J | N | C | 9/29/21 | |
| 7n8h | C | B | A | 7/14/21 | |
| 6wo3 | H | L | E | 8/19/20 | |
| 6wmw | M | N | B | 7/15/20 | |
| 6wmw | H | L | B | 7/15/20 | |
| 7czx | I | M | B | 3/10/21 | |
| 7czw | H | L | A | 3/10/21 | |
| 7rah | B | A | E | 9/15/21 | |
| 7rah | D | C | E | 9/15/21 | |
| 7kn4 | M | N | B | 9/22/21 | |
| 6nyq | H | L | C | 1/22/20 | |
| 6xkr | H | L | P | 9/9/20 | yes |
| 6jjp | D | E | F | 10/30/19 | |
| 7lxy | N | O | J | 4/14/21 | |
| 7lxy | E | D | A | 4/14/21 | |
| 7lr4 | H | L | D | 12/15/21 | |
| 7r89 | C | D | BA | 9/8/21 | |
| 6ywc | D | E | F | 10/7/20 | |
| 6xxv | D | E | F | 4/22/20 | |
| 7ly2 | N | O | J | 4/14/21 | |
| 6vvu | G | I | D | 12/30/20 | |
| 6k0y | A | B | C | 12/11/19 | yes |
| 6v4o | H | L | N | 10/7/20 | |
| 6wgl | A | B | C | 9/16/20 | |
| 6xsw | D | E | F | 7/21/21 | |
| 6hx4 | H | L | B | 10/30/19 | |
| 6wzk | A | B | E | 11/25/20 | |
| 7lxz | H | J | A | 4/14/21 | |
| 7ceb | C | D | B | 6/23/21 | |
| 6lxi | C | D | B | 12/2/20 | |
| 6osv | H | L | K | 4/1/20 | |

| | | | | | | |
|---|---|---|---|---|---|---|
| 7lbg | H | G | A | 3/10/21 | | |
| 7lbg | F | E | A | 3/10/21 | | yes |
| 6a3w | A | B | C | 10/10/18 | | |
| 7bnv | H | L | A | 11/17/21 | yes | yes |
| 7lm9 | H | L | A | 3/31/21 | | |
| 7o52 | H | L | U | 7/28/21 | | |
| 7s4s | H | L | A | 9/22/21 | | |
| 7lop | X | Y | Z | 3/3/21 | | |
| 6xkp | M | N | B | 10/14/20 | | |
| 6h2y | H | L | D | 8/14/19 | | |
| 6phc | C | D | E | 10/2/19 | | yes |
| 6osh | H | L | K | 4/1/20 | | |
| 7n3d | H | L | C | 7/7/21 | | |
| 7e7y | A | B | R | 6/9/21 | | |
| 7chf | A | B | R | 9/16/20 | | yes |
| 7lm8 | H | L | A | 3/31/21 | | yes |
| 7kn3 | M | N | B | 9/22/21 | | yes |
| 7n0u | H | L | C | 8/11/21 | | |
| 7joo | H | L | C | 10/14/20 | | |
| 7dk2 | D | E | F | 12/8/21 | | yes |
| 6sni | H | L | X | 3/11/20 | | |
| 7c88 | A | B | C | 4/14/21 | | yes |
| 7dc8 | B | A | C | 1/13/21 | | |
| 7mmo | D | E | F | 5/12/21 | | |
| 7kfv | F | G | E | 12/2/20 | | yes |
| 7k9z | B | A | E | 10/28/20 | | |
| 6mlk | H | L | A | 10/17/18 | | |
| 6mg7 | H | L | G | 9/25/19 | | yes |
| 7l0l | H | L | BA | 11/3/21 | yes | |
| 6ogx | C | D | G | 7/10/19 | | |
| 7lcv | A | B | C | 6/9/21 | | |
| 6pi7 | F | E | D | 7/24/19 | | |
| 7ps6 | H | L | E | 12/15/21 | yes | yes |
| 6rlo | G | H | L | 5/12/21 | | |
| 7lab | Y | X | B | 3/10/21 | | |
| 7cm4 | H | L | A | 1/20/21 | | |
| 6gv4 | H | L | BA | 11/21/18 | | |
| 6j14 | A | B | G | 11/6/19 | | |
| 6ohg | C | B | A | 6/17/20 | | |
| 6v4n | D | E | W | 10/7/20 | | |

| | | | | | | |
|---|---|---|---|---|---|---|
| 7e5o | H | L | A | 9/8/21 | | yes |
| 6p67 | E | F | G | 9/4/19 | | |
| 7ean | H | L | A | 3/31/21 | | yes |
| 7eam | H | L | A | 3/17/21 | | |
| 6ppg | B | A | G | 12/11/19 | | yes |
| 6nmr | J | K | M | 8/7/19 | | |
| 7djz | A | B | C | 6/9/21 | | yes |
| 6m3b | C | B | A | 7/8/20 | | yes |
| 6o1f | H | L | AI | 10/16/19 | | |
| 7lr3 | H | L | D | 12/15/21 | | |
| 6r8x | C | B | A | 4/10/19 | | yes |
| 6wxl | F | E | DC | 6/9/21 | | |
| 6a67 | H | L | A | 8/29/18 | | |
| 6mej | H | L | C | 11/21/18 | | yes |
| 6mej | A | B | C | 11/21/18 | | |
| 6udj | H | I | J | 1/29/20 | | |
| 7cho | B | C | A | 5/19/21 | | yes |
| 7bwj | H | L | E | 6/3/20 | | yes |
| 7kqb | H | L | A | 5/26/21 | | |
| 6kz0 | K | L | J | 5/27/20 | | |
| 7vux | H | L | A | 11/17/21 | yes | |
| 7q0i | H | L | C | 12/22/21 | yes | yes |
| 6pzw | H | L | A | 12/4/19 | | |
| 7orb | E | F | X | 7/7/21 | | |
| 6nmu | B | A | C | 8/7/19 | | |
| 7s0b | C | D | E | 10/6/21 | | |
| 6rps | H | L | A | 11/13/19 | | |
| 7l7e | O | P | K | 9/1/21 | | |
| 7orb | H | L | R | 7/7/21 | | |
| 7r6w | A | B | R | 7/21/21 | | |
| 7r6w | H | L | R | 7/21/21 | | |
| 7kyl | H | L | E | 4/7/21 | | |
| 6iut | H | L | A | 1/16/19 | | |
| 6nmt | B | A | C | 8/7/19 | | |
| 6dkj | A | B | D | 5/8/19 | | yes |
| 7c61 | H | L | A | 7/29/20 | | |
| 7m3n | H | L | A | 7/28/21 | | |
| 7s13 | H | L | D | 10/20/21 | | |
| 7m7w | C | D | S | 5/5/21 | | |
| 6nmv | H | L | S | 8/7/19 | | yes |

| | | | | | | |
|---|---|---|---|---|---|---|
| 6icc | H | L | A | 2/13/19 | | |
| 7m7w | H | L | R | 5/5/21 | | yes |
| 6j15 | A | B | C | 11/6/19 | | yes |
| 6whk | C | B | A | 4/14/21 | | |
| 7kpb | H | L | AC | 1/13/21 | | |
| 6pe8 | H | L | T | 8/14/19 | | |
| 7ket | A | B | C | 6/9/21 | | |
| 6ion | H | L | A | 1/15/20 | | yes |
| 6nms | B | A | C | 8/7/19 | | |
| 6lz9 | H | L | B | 3/11/20 | | yes |
| 6p3r | C | D | E | 5/27/20 | | |
| 7coe | B | C | D | 8/4/21 | | |
| 7ps4 | H | L | E | 12/15/21 | yes | |
| 7ps1 | A | B | E | 12/15/21 | | yes |
| 6phb | D | C | I | 10/2/19 | | |
| 6lgw | C | D | F | 2/19/20 | | |
| 7s11 | I | M | D | 11/3/21 | yes | |
| 6ieb | E | F | B | 4/10/19 | | |
| 6e63 | H | L | P | 11/28/18 | | yes |
| 7seg | H | L | C | 11/24/21 | | yes |
| 6wh9 | E | F | D | 9/9/20 | | |
| 7ps0 | H | L | E | 12/15/21 | yes | yes |
| 6xlq | B | C | A | 9/2/20 | | |
| 7dm1 | D | C | A | 12/23/20 | | |
| 7dm2 | H | L | A | 12/23/20 | | yes |
| 6j5d | H | L | A | 2/6/19 | | |
| 7o9s | H | L | A | 6/23/21 | | yes |
| 7kmg | D | E | F | 1/27/21 | | |
| 6q0e | H | L | A | 12/18/19 | | |
| 7cr5 | H | L | A | 3/24/21 | | |
| 7q0g | A | B | E | 12/22/21 | yes | yes |
| 6h3t | I | M | B | 2/27/19 | | |
| 6s5a | H | L | DA | 9/25/19 | | |
| 7dha | C | B | A | 9/22/21 | | |
| 6ddm | B | A | C | 10/24/18 | | |
| 7mzm | H | L | A | 10/6/21 | yes | yes |
| 6nha | H | L | AB | 12/25/19 | | |
| 6xpx | B | C | A | 5/19/21 | | |
| 6xq0 | E | F | D | 5/19/21 | | |
| 6e56 | H | J | A | 5/22/19 | | |

| | | | | | | |
|---|---|---|---|---|---|---|
| 7ps2 | H | L | G | 12/15/21 | yes | yes |
| 7ps2 | A | B | G | 12/15/21 | | yes |
| 6tyb | H | L | G | 10/2/19 | | |
| 6u9s | D | E | F | 5/13/20 | | yes |
| 7bq5 | H | L | A | 3/24/21 | | |
| 7dr4 | A | B | J | 4/14/21 | | yes |
| 6m58 | C | D | B | 4/29/20 | | |
| 6ss2 | H | L | A | 6/10/20 | | |
| 7kzb | H | L | C | 2/3/21 | | yes |
| 6vug | D | C | B | 2/17/21 | | |
| 6e4x | Z | Y | B | 5/22/19 | | yes |
| 6otc | H | L | A | 6/5/19 | | |
| 6oy4 | D | C | A | 8/28/19 | | yes |
| 7nx8 | H | L | E | 4/7/21 | | yes |
| 7bel | C | D | X | 3/3/21 | | |
| 7mzg | H | L | A | 10/6/21 | | |
| 7bek | H | L | E | 3/3/21 | | |
| 7mf1 | H | L | A | 5/12/21 | | |
| 7bei | H | L | E | 3/3/21 | | yes |
| 7bel | E | F | X | 3/3/21 | | |
| 6iuv | C | D | B | 1/16/19 | | yes |
| 6iea | H | L | A | 4/10/19 | | |
| 7e3o | H | L | R | 9/15/21 | | |
| 7ahu | B | A | EF | 7/7/21 | | |
| 6q18 | H | L | A | 12/18/19 | | |
| 7kmh | A | B | C | 1/27/21 | | |
| 7mzk | N | M | B | 10/6/21 | yes | |
| 7or9 | H | L | E | 7/7/21 | | yes |
| 6oz2 | H | L | G | 8/19/20 | | |
| 6xzw | H | L | D | 10/14/20 | | yes |
| 6iek | B | C | A | 4/10/19 | | yes |
| 7rks | I | M | S | 9/22/21 | | |
| 6w7s | H | L | A | 9/9/20 | | yes |
| 7mzi | H | L | A | 10/6/21 | | |
| 7neh | H | L | E | 3/3/21 | | |
| 7mzj | H | L | A | 10/6/21 | yes | yes |
| 6i8s | E | I | A | 2/13/19 | | |
| 6oor | H | L | A | 7/17/19 | | |
| 6vy6 | H | L | A | 1/6/21 | | yes |
| 7bbj | H | L | A | 12/29/21 | yes | |

| | | | | | | |
|---|---|---|---|---|---|---|
| 6mfp | C | D | A | 9/18/19 | | |
| 6wm9 | E | F | D | 1/27/21 | | |
| 7mzh | H | L | E | 10/6/21 | | |
| 7bep | A | B | E | 3/3/21 | | |
| 7chz | H | L | I | 1/13/21 | | yes |
| 6qig | H | L | A | 9/4/19 | | |
| 6hhc | H | L | A | 9/11/19 | | yes |
| 6woz | K | L | J | 1/27/21 | | |
| 7lsg | H | L | C | 4/7/21 | | yes |
| 7jx3 | C | D | R | 10/14/20 | | |
| 7jx3 | H | L | R | 10/14/20 | | yes |
| 6j6y | E | F | D | 8/7/19 | | |
| 7kq7 | H | L | B | 4/7/21 | | |
| 6ztr | A | B | J | 5/5/21 | | yes |
| 7ce2 | Z | B | A | 4/7/21 | | |
| 6wtu | E | F | D | 1/27/21 | | |
| 6ocb | H | L | A | 5/29/19 | | |
| 6wds | H | L | CAB | 7/15/20 | | |
| 7kyo | H | L | B | 8/25/21 | | |
| 6o39 | B | A | C | 4/3/19 | | |
| 6ba5 | F | E | O | 6/13/18 | | |
| 6n6b | K | L | A | 7/3/19 | | |
| 6ye3 | G | H | I | 12/30/20 | | yes |
| 7bsc | H | L | A | 12/23/20 | | |
| 6kyz | B | C | A | 5/27/20 | | |
| 7ec5 | E | F | BAC | 3/31/21 | | |
| 6mhr | A | B | C | 11/21/18 | | |
| 6pzf | F | E | B | 12/4/19 | | |
| 6pze | H | L | A | 12/4/19 | | |
| 6z2l | C | B | A | 7/22/20 | | |
| 6e3h | H | L | BA | 9/26/18 | | |
| 6cxy | H | L | C | 4/10/19 | | |
| 6nz7 | H | L | BA | 5/8/19 | | |
| 6vy4 | C | D | B | 12/30/20 | | |
| 7e72 | C | D | F | 11/10/21 | yes | yes |
| 7n4j | H | L | A | 10/6/21 | yes | |
| 6e62 | H | L | P | 11/28/18 | | |
| 6q20 | H | L | A | 10/23/19 | | |
| 6vc9 | H | L | A | 11/11/20 | | |
| 6lyn | H | L | D | 2/24/21 | | |

| | | | | | |
|---|---|---|---|---|---|
| 6ivz | H | L | A | 2/13/19 | |
| 6id4 | C | D | EF | 2/6/19 | |
| 6hxw | C | D | B | 8/28/19 | |
| 7d85 | E | F | D | 4/7/21 | |
| 7r8l | H | L | E | 8/4/21 | |
| 7klh | H | L | A | 2/10/21 | |
| 7mhy | O | P | A | 6/16/21 | yes |
| 7mhy | M | N | A | 6/16/21 | |
| 6hga | H | L | B | 3/18/20 | |
| 6pxh | H | L | B | 9/25/19 | |
| 7cj2 | K | L | B | 7/14/21 | |
| 6j11 | F | G | B | 7/24/19 | |
| 7jie | E | F | A | 6/30/21 | |
| 6n5e | G | F | B | 6/5/19 | |
| 6u6u | H | L | R | 4/22/20 | |
| 6iap | E | D | A | 6/12/19 | |
| 6iap | H | L | A | 6/12/19 | yes |
| 7n3c | H | L | C | 7/7/21 | |
| 7e9b | H | L | C | 7/28/21 | |
| 7kpg | H | L | S | 12/16/20 | |
| 6jep | H | L | E | 5/15/19 | |
| 6dfj | H | L | E | 10/24/18 | yes |
| 6ute | C | D | S | 4/15/20 | yes |
| 6ddr | B | A | C | 10/24/18 | yes |
| 6ddv | B | A | C | 10/24/18 | yes |
| 6a77 | H | L | A | 1/30/19 | |
| 6rvc | D | | A | 10/2/19 | |
| 6gju | C | | A | 6/26/19 | |
| 6gjq | B | | A | 6/19/19 | |
| 6r7t | A | | B | 5/1/19 | |
| 7l1v | S | | R | 2/10/21 | |
| 7my3 | E | | A | 6/16/21 | |
| 7kkl | E | | C | 11/11/20 | |
| 6v7y | F | | A | 9/16/20 | |
| 6x04 | H | | G | 12/9/20 | yes |
| 7rnn | C | | D | 8/11/21 | |
| 7p77 | A | | B | 8/4/21 | |
| 6u54 | A | | B | 11/6/19 | |
| 7mjh | F | | C | 5/12/21 | |
| 7my2 | H | | E | 6/16/21 | |

| | | | | | |
|---|---|---|---|---|---|
| 7kn6 | C | A | 1/20/21 | | yes |
| 7kjh | A | C | 2/3/21 | | |
| 7o06 | A | C | 9/8/21 | | yes |
| 6zxn | E | B | 9/23/20 | | |
| 7a29 | E | B | 10/21/20 | | |
| 6ze1 | B | A | 6/30/21 | | yes |
| 7d6y | B | A | 10/6/21 | | |
| 6rnk | B | A | 8/14/19 | | |
| 6v7z | F | AB | 9/16/20 | | |
| 7kn5 | E | A | 1/20/21 | | |
| 7kn5 | C | A | 1/20/21 | | yes |
| 6u55 | A | B | 11/6/19 | | |
| 6yu8 | B | A | 2/17/21 | | |
| 7apj | B | A | 8/25/21 | | |
| 6x05 | K | A | 12/9/20 | | |
| 7olz | B | A | 8/11/21 | | |
| 7olz | C | A | 8/11/21 | | |
| 6qup | B | A | 8/5/20 | | yes |
| 6qgw | B | A | 6/26/19 | | |
| 6z20 | D | C | 9/23/20 | | |
| 6rqm | B | A | 7/8/20 | | |
| 6oyh | E | A | 7/10/19 | | |
| 6os2 | D | A | 2/19/20 | | yes |
| 6qgx | B | A | 6/26/19 | | |
| 6qgy | B | A | 6/26/19 | | |
| 6z1v | B | A | 9/23/20 | | |
| 7o0s | A | B | 9/15/21 | | |
| 7r98 | F | C | 7/7/21 | | |
| 7c8v | A | B | 6/24/20 | | |
| 6ir1 | B | A | 11/13/19 | | yes |
| 7cz0 | E | A | 9/8/21 | | |
| 6x07 | B | A | 12/9/20 | | yes |
| 7nfq | C | A | 12/1/21 | yes | |
| 7nfr | B | A | 12/1/21 | yes | |
| 6lz2 | B | A | 12/23/20 | | |
| 6gs4 | H | A | 1/30/19 | | |
| 6gk4 | F | D | 6/19/19 | | |
| 6o3c | B | A | 7/3/19 | | |
| 7nx0 | D | C | 10/27/21 | yes | |
| 6yz5 | F | E | 6/3/20 | | |

| | | | | | |
|---|---|---|---|---|---|
| 6z6v | G | BC | 6/10/20 | | |
| 6rtw | B | A | 10/2/19 | | |
| 7k84 | B | A | 10/14/20 | | |
| 6uc6 | D | B | 3/4/20 | | |
| 7lzp | G | A | 12/22/21 | | yes |
| 6oq6 | D | A | 7/10/19 | | |
| 6ir2 | B | A | 11/13/19 | | yes |
| 6gkd | C | A | 6/19/19 | | |
| 7d30 | A | B | 2/17/21 | | |
| 7lzp | F | D | 12/22/21 | | yes |
| 7lzp | E | D | 12/22/21 | yes | |
| 6mxt | N | A | 11/14/18 | | |
| 6dbg | C | B | 7/18/18 | | |
| 6vbg | D | B | 11/25/20 | | |
| 7azb | B | A | 11/25/20 | | |
| 6gwn | B | A | 1/1/20 | | |
| 6gwn | C | A | 1/1/20 | | |
| 6zg3 | E | AI | 3/3/21 | | |
| 7a0v | B | A | 12/30/20 | | |
| 6ssi | J | E | 2/12/20 | | |
| 6gjs | C | A | 6/26/19 | | |
| 6gjs | B | A | 6/26/19 | | |
| 6lr7 | B | A | 4/29/20 | | yes |
| 7vnb | A | B | 11/24/21 | yes | |
| 7ldj | G | C | 5/5/21 | | yes |
| 7anq | B | A | 10/20/21 | yes | yes |
| 6h02 | B | A | 8/29/18 | | |
| 6hhu | G | A | 7/24/19 | | yes |
| 6hhu | H | A | 7/24/19 | | |
| 6uft | B | A | 3/4/20 | | yes |
| 7e53 | B | A | 10/13/21 | yes | |
| 6oca | C | A | 4/1/20 | | |
| 7m1h | G | A | 12/22/21 | yes | |
| 7m1h | E | A | 12/22/21 | yes | |
| 7m1h | F | A | 12/22/21 | yes | |
| 7kdu | C | BA | 8/4/21 | | |
| 6fyu | C | BA | 11/7/18 | | |
| 6h6y | G | C | 12/19/18 | | |
| 7na9 | D | A | 12/22/21 | yes | yes |
| 6ui1 | D | A | 3/4/20 | | yes |

| | | | | | |
|---|---|---|---|---|---|
| 7t5f | E | D | 12/29/21 | yes | |
| 7kc9 | F | ED | 8/4/21 | | |
| 7lvw | I | D | 3/24/21 | | |
| 7aqy | C | B | 11/3/21 | yes | |
| 6zrv | B | A | 8/26/20 | | |
| 7t5f | C | A | 12/29/21 | yes | yes |
| 7kbk | C | AB | 8/4/21 | | |
| 7kd2 | C | BA | 8/4/21 | | |
| 6ul6 | C | A | 3/4/20 | | |
| 6ul6 | B | A | 3/4/20 | | |
| 6i8g | B | A | 10/2/19 | | |
| 6h16 | B | A | 1/30/19 | | |
| 7kd0 | C | BA | 8/4/21 | | |
| 7aqz | D | A | 11/3/21 | | |
| 7l6v | B | A | 12/22/21 | yes | |
| 7l6v | D | A | 12/22/21 | yes | |
| 7l6v | C | A | 12/22/21 | yes | |
| 7l6v | F | A | 12/22/21 | yes | |
| 6uht | C | A | 3/4/20 | | |
| 7n0r | D | B | 6/2/21 | | |
| 6oq7 | C | A | 7/10/19 | | |
| 6rah | C | B | 7/31/19 | | |
| 7ar0 | B | A | 11/3/21 | yes | |
| 6xw4 | C | A | 4/22/20 | | |
| 6h15 | D | B | 1/30/19 | | |
| 6h72 | C | A | 12/19/18 | | |
| 7aqx | D | B | 11/3/21 | | |
| 6sge | B | C | 9/25/19 | | |
| 6waq | A | B | 4/1/20 | | |
| 7n0i | L | GH | 6/9/21 | | |
| 7d2z | A | B | 2/17/21 | | |
| 6ocd | B | A | 4/1/20 | | |
| 6tej | C | B | 4/1/20 | | |
| 6oq8 | D | A | 7/10/19 | | |
| 6b20 | F | A | 5/30/18 | | |
| 6obe | B | A | 4/1/20 | | |
| 6obc | B | A | 4/1/20 | | |
| 6obo | C | A | 4/1/20 | | yes |
| 6i6j | C | A | 2/27/19 | | yes |
| 7mfu | B | A | 6/2/21 | | yes |

| | | | | |
|---|---|---|---|---|
| 7mfu | F | D | 6/2/21 | |
| 7kgj | B | A | 2/3/21 | yes |
| 7kgk | B | A | 2/3/21 | |
| 7now | C | D | 4/7/21 | yes |
| 7nqa | D | A | 7/21/21 | |
| 6xzu | A | B | 8/12/20 | |
| 7czd | A | B | 7/14/21 | |

### References


1 Stranges, P. B. & Kuhlman, B. A comparison of successful and failed protein interface designs highlights the challenges of designing buried hydrogen bonds. *Protein Sci* **22**, 74-82, doi:10.1002/pro.2187 (2013).

2 Alford, R. F. *et al.* The Rosetta All-Atom Energy Function for Macromolecular Modeling and Design. *J Chem Theory Comput* **13**, 3031-3048, doi:10.1021/acs.jctc.7b00125 (2017).

3 Fu, L., Niu, B., Zhu, Z., Wu, S. & Li, W. CD-HIT: accelerated for clustering the next-generation sequencing data. *Bioinformatics* **28**, 3150-3152, doi:10.1093/bioinformatics/bts565 (2012).

4 Robin, X. *et al.* pROC: an open-source package for R and S+ to analyze and compare ROC curves. *BMC Bioinformatics* **12**, 77, doi:10.1186/1471-2105-12-77 (2011).

5 Wei, R. & Wang, J. multiROC: Calculating and Visualizing ROC and PR Curves Across Multi-Class assifications. (2018).

6 Pierce, B. G., Hourai, Y. & Weng, Z. Accelerating protein docking in ZDOCK using an advanced 3D convolution library. *PLoS One* **6**, e24657, doi:10.1371/journal.pone.0024657 (2011).

7 Vreven, T., Hwang, H. & Weng, Z. Integrating atom-based and residue-based scoring functions for protein-protein docking. *Protein Sci* **20**, 1576-1586, doi:10.1002/pro.687 (2011).
